# Mechanism of dopamine binding and allosteric modulation of the human D1 dopamine receptor

**DOI:** 10.1101/2021.02.07.430101

**Authors:** Youwen Zhuang, Brian Krumm, Huibing Zhang, X. Edward Zhou, Yue Wang, Xi-Ping Huang, Yongfeng Liu, Xi Cheng, Yi Jiang, Hualiang Jiang, Cheng Zhang, Wei Yi, Bryan L. Roth, Yan Zhang, H. Eric. Xu

## Abstract

Dopamine is an essential neurotransmitter, which functions are mediated by five G protein-coupled receptors, dopamine D1 to D5 receptors (D1R-D5R) in mammals. Among them, D1R is the most abundantly expressed dopamine receptor in the CNS and is the central receptor mediating excitatory dopamine signaling in multiple dopaminergic pathways. Dysregulation of D1R signaling has been directly linked to Parkinson’s disease (PD), schizophrenia, and drug abuse. Due to its fundamental functions in human diseases, D1R has long been the subject of intensive drug development effort toward the treatment of neuropsychiatric diseases. Here, we report the structures of D1R-Gs complex bound to endogenous agonist dopamine and synthetic agonist SKF81297, both with positive allosteric modulator LY3154207. These structures reveal the basis of dopamine recognition, the binding and potential allosteric regulation of DRD1 PAM LY3154207, and provide structural templates for design of subtype-selective D1R ligand for drug discovery targeting DRD1 for treating various CNS diseases.

---

Dopamine acts as an essential neurotransmitter whose signaling is conducted through five G protein-coupled receptors (GPCRs), dopamine D1 to D5 receptors (DRD1-DRD5)^1^. The D1-like receptors, comprising DRD1 and DRD5, primarily couple to the G_s_ family of G proteins to activate adenylyl cyclase and induce cAMP production. DRD1 is the most abundantly expressed dopamine receptor in the CNS^1^. It is the central receptor mediating excitatory dopamine signaling in multiple dopaminergic pathways. Dysregulation of DRD1 signaling has been directly linked to Parkinson’s disease (PD), schizophrenia, and drug abuse^1,2^. Due to its fundamental functions in human diseases, DRD1 has long been the subject of intensive drug development efforts toward the treatment of neuropsychiatric diseases^3^. A majority of DRD1 agonists, including the SKF compounds, targets the orthosteric pocket of DRD1, but none has passed clinical trials for neuropsychiatric symptoms to date^3^.

GPCR positive allosteric modulators (PAMs) have been proposed to provide unique advantages over orthosteric agonists including greater receptor subtype selectivity, saturable therapeutic effects and the ability to maintain spatial and temporal patterns of endogenous dopamine signaling, which collectively may lead to reduced side effects^3,4^. Multiple groups have reported DRD1 positive allosteric modulators (PAMs), such as LY3154207, CID2886111, and DETQ, to stimulate DRD1 signaling^5,6^ With ongoing clinical investigation, DRD1 PAMs may offer new therapeutic opportunities for PD^3,7^.

Despite significant efforts, the structural basis of DRD1 ligand binding and allosteric regulation properties remains poorly understood, which has significantly impeded the discovery of potential DRD1-selective drugs with minimal side effects. Here we report two structures of DRD1-G_s_ complexes activated by the endogenous ligand, dopamine, and a synthetic agonist, SKF81297, both in the presence of LY3154207, respectively (Figs. 1a and 1b). We used an engineered miniG_s_ (miniG_αs__DN) to assembly DRD1-G_s_ signaling complexes ^8^ (Supplementary information, Fig. S1). To obtain stable DRD1-G_s_ complexes for structural studies, we co-expressed the wild-type human DRD1, miniG_αs__DN, rat G_β1_ and bovine G_γ2_ in Sf9 insect cells. The complexes were prepared as described in the Methods section and purified to homogeneity for single particle cryo-EM studies (Supplementary information, Fig. S2). Two different DRD1 PAMs, CID2886111 and LY3154207, which bind to different sites on DRD1 as shown by prior studies^5,6^, were added to further stabilized the dopamine-bound DRD1-G_s_ complex. The structures of DRD1-G_s_ complexes with dopamine/LY3154207 and SKF81297/LY3154207 were determined at a global resolution of 3.2 Å and 3.0 Å, respectively (Figs. 1a and 1b; Supplementary information, Fig. S3 and Table S1). The relatively high-resolution maps allowed us to unambiguously model most portions of DRD1 from S21 to Y348, the G_s_ heterotrimer, the orthosteric agonists, and the nanobody Nb35 (Supplementary information, Figs. S4 and S5a). In addition, in the SKF81297-bound DRD1 structure, clear density for the PAM LY3154207 was observed above ICL2 (Fig. 1b; Supplementary information, Figs. S4 and S5a), allowing us to define the binding pose of LY3154207 and the allosteric site. In the dopamine-bound DRD1 structure, the binding pose of LY3154207 can be defined (Fig. 1b; Supplementary information, Figs. S4 and S5a), but no density was observed for CID2886111.

**Fig. 1.**
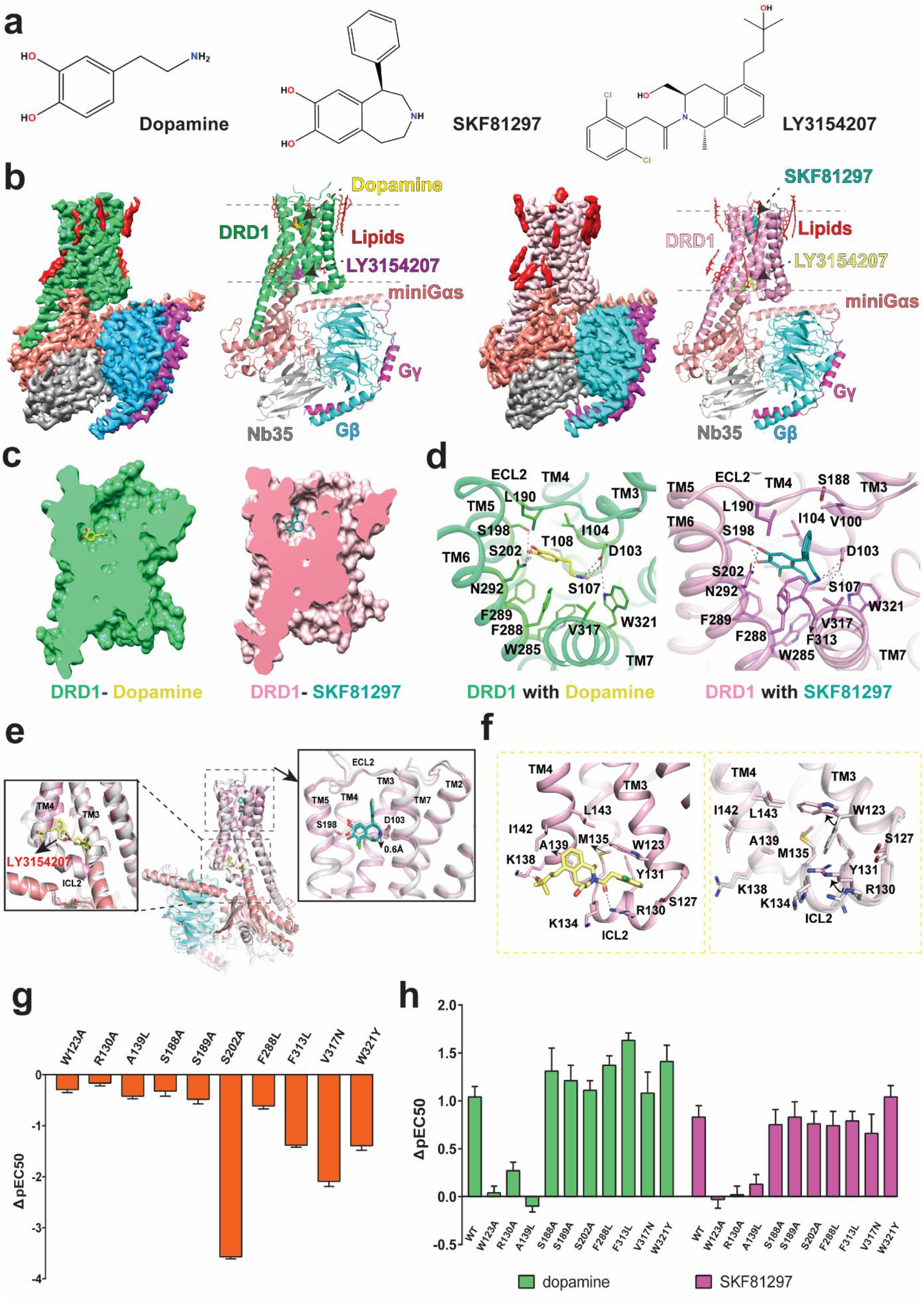
Structures of DRD1-G_s_ complexes. **a** Chemical structures of ligands. **b** EM maps and structures of the DRD1-dopamine-G_s_ complex and the DRD1-SKF81297-G_s_ complex, both with LY3154207. The maps are shown at 0.045 and 0.083 threshold for DRD1-dopamine/LY3154207-G_s_ complex and DRD1-SKF81297/LY3154207-G_s_ complex, respectively. **c** The dopamine and SKF81297 orthosteric binding pockets. **d** Interactions of dopamine and SKF81297 with DRD1. **e** Superposition of DRD1-SKF81297-G_s_ structures with or without LY3154207. The structure without PAM: white; The structure with PAM, DRD1: pink, SKF81297: teal, LY3154207: yellow orange, G_αs_: salmon, G_β_: cyan, G_γ_: magenta. The hydrogen bond interactions are shown as black dashed line and yellow dashed line for DRD1-SKF81297 structure and DRD1-SKF81297/LY3154207 structure, respectively. **f** The binding mode of LY3154207 and the conformational differences of the DRD1 allosteric binding site with or without LY3154207. The LY3154207 was removed for better presentation of the alignment. **g** cAMP accumulation analysis of WT DRD1 and DRD1 mutants activated by dopamine. Data are presented as mean values ± SEM with a minimum of two technical replicates and n = 3 biological replicates. Greek letter delta (Δ) for the difference (ΔpEC50) when compared with the wild-type receptor values. **h** Comparison of dopamine and SKF81297 in cAMP accumulation assays with WT and mutated DRD1 with or without 30 nM LY3154207. Data are presented as mean values ± SEM with a minimum of two technical replicates and n = 3 biological replicates. Greek letter delta (Δ) for the difference of pEC50 values (ΔpEC50) when comparing the pEC50s of WT or each mutant with and without 30 nM LY3154207.

The overall structures of LY3154207-bound DRD1 with dopamine and SKF81297 are quite similar, with a root mean square deviation (RMSD) value of 0.6 Å for the main chain Cα atoms. However, the orthosteric binding pocket of SKF81297 is narrower compared to that of dopamine (Fig. 1c; Supplementary information, Fig. S5b). In both structures, DRD1 adopts a canonical seven-helical transmembrane domain (TMD), the ligand binding pockets are located at the extracellular part of the TMD and the G-protein coupling interface is located at the cytoplasmic side (Figs. 1a and 1b).

In the dopamine-bound DRD1 structure, dopamine occupies the orthosteric binding pocket (OBP) composed of residues from TM3, TM5-7 and capped by extracellular loop 2 (ECL2) (Fig. 1d; Supplementary information, Fig. S5a). The primary amine group forms direct ionic contacts with the carboxylate group of D103^3.32^ (superscript based on Ballesteros-Weinstein numbering rules of GPCRs), which is highly conserved in aminergic GPCRs^9^. Such interaction is further enhanced by hydrogen bond interactions among D103^3.32^, S107^3.36^, and W321^7.43^ (Fig. 1d). The catechol moiety forms hydrogen bond interactions with S198^5.42^ and S202^5.46^ from TM5 and N292^6.55^ from TM6 (Fig. 1d). These findings agree well with the mutational results from previous studies reporting that S198^5.42^ and S202^5.46^ are pivotal for dopamine binding^10^. In addition to the polar interaction network, hydrophobic residues I104^3.33^, L190^ECL2^, W285^6.48^, F288^6.51^, F289^6.52^ and V317^7.39^ form extensive hydrophobic interactions with dopamine to further stabilize the dopamine binding (Fig. 1d). For SKF81297, although it shares the same catechol moiety as dopamine and its benzazepine ring overlaps well with the phenylethylamine moiety of dopamine, the conformation of the catechol group of SKF81297 is slightly different from that of dopamine (Fig. 1d; Supplementary information, Fig. S5c). As a result, the catechol moiety of SKF81297 forms hydrogen bonds with S198^5.42^ but not S202^5.46^ (Supplementary information, Fig. S5c). The extra benzene group of SKF81297 occupies a small extended binding pocket (EBP) at the extracellular vestibule formed by residues V100^3.29^, L190^ECL2^, S198^5.42^ and F313^7.35^ (Supplementary information, Fig. S5c), which contribute to its higher affinity to DRD1 than that of dopamine. Interestingly, the side chain of D187^ECL2^ points towards polar residues K81^2.60^ and D314^7.36^ in the SKF81297-bound DRD1 but not in the dopamine-bound DRD1, forming a potential polar interaction network (Supplementary information, Fig. S5d). The clustering of the side chains of these three polar residues leads to a narrower ligand-binding pocket for SKF81297 than that for dopamine.

To validate the structural findings in dopamine binding pockets in DRD1, we mutated residues near the pockets and analyzed the expression levels and cAMP accumulation effects of these DRD1 mutants when activated by dopamine. Corresponding to the binding modes, mutations of residues D103^3.32^, S198^5.42^ and N292^6.55^ largely decrease the potency of dopamine (Supplementary information, Fig. S6 and Table S2). Furthermore, mutations of nearby residues including K81^2.61^, I104^3.33^, S107^3.36^, L190^ECL2^, S199^5.43^, F288^6.51^ and W321^7.43^ also decreased dopamine potency (Supplementary information, Figs. S6 and S7; Tables S2 and S3), supporting the binding mode of dopamine to DRD1.

DRD1 has been proposed to possess at least two different positive allosteric sites, one of these sites has been well characterized for several potent DRD1 PAMs, including DETQ and LY3154207 based on computational simulations and extensive mutagenesis data^5,6^. In our structure, the contact pattern of LY3154207 with DRD1 is quite different from that in a previously reported simulation model of LY3154207-bound DRD1^6^. The whole LY3154207 molecule lies in the cleft between TM3 and TM4 and right above ICL2 with a boat conformation, which is about 33 Å away from the orthosteric DRD1 pocket when measured at the Cα atoms of D103^3.32^ and Y131^ECL2^ (Fig. 1e). A similar allosteric site has also been identified in the β_2_-adrenergic receptor (β_2_AR) for a β_2_AR PAM named Cmpd-6FA^11^ (Supplementary information, Fig. S8). In the allosteric site, LY3154207 mainly forms hydrophobic and van der Waals interactions with DRD1 (Fig. 1f), which is consistent with the hydrophobic property of LY3154207. The dichlorophenyl group of LY3154207 is sandwiched by the side chains of R130^ICL2^ and W123^3.52^ to form cation-*π* and _π-π_ interactions, respectively (Fig. 1f). The central tetrahydroisoquinoline (THIQ) ring of LY3154207 forms hydrophobic interactions with surrounding residues M135^ICL2^, A139^4.41^, I142^4.44^ and L143^4.45^. In addition, hydrogen bonds between LY3154207 and polar residues R130^ICL2^, K134^ICL2^ and K138^4.40^ are also observed (Fig.1f).

To correlate the function and binding mode of LY3154207, we firstly analyzed the effects of DRD1 orthosteric site mutations on LY3154207 efficacy and potency. The G_s_-mediated cAMP accumulation results indicated that most of the mutations have minimal effects on LY3154207 binding, including residues D103^3.32^, S198^5.42^, S199^5.43^ and F288^6.51^ (Supplementary information, Fig. S6 and Table S2), which were important for dopamine and SKF81297 binding. The addition of LY3154207 increases cAMP accumulation efficacy and the potency of both dopamine and SKF81297 in WT DRD1 and DRD1 orthosteric pocket mutants (Fig. 1g and 1h; Supplementary information, Fig. S7 and Table S3), suggesting that orthosteric agonist and LY3154207 conduct cooperative effects on G_s_ stimulation. Subsequently, we mutated most residues around the LY3154207 pocket and tested the abilities of G protein recruitment of DRD1 allosteric site mutants. The presence of LY3154207 increases potency of dopamine and SKF81297 by about one Log (Figs. 1g and 1h). Mutations of W123A, R130A, and A139L in the allosteric binding site nearly abolished the allosteric effects of LY3154207 on DRD1 activation potency of dopamine and SKF81297, while these mutations had modest effects on the function of ligand binding to the orthosteric site (Figs. 1g and 1h; Supplementary information, Figs. S7 and S9; Tables S3 and S4). Interestingly, LY3154207 alone can activate DRD1 to a certain extent (Supplementary information, Fig. S6 and Table S2).

LY3154207 shares a high chemical similarity with DETQ. The only difference is that LY3154207 contains a longer alkyl linker between the C5 tertiary alcohol and the THIQ ring (Supplementary information, Fig. S10a). It is likely that DETQ occupies the same allosteric site as LY3154207. Supporting this hypothesis, previous studies indicated that residues W123^3.52^, R130^ICL2^ and L143^4.45^ were crucial for DETQ potency, which all directly interact with LY3154207 in the allosteric site^5^. Also, A139^4.41^ has been shown to play a key role in the selectivity of DETQ for DRD1 over D5R^5^. In our structure, A139^4.41^ in TM4 forms hydrophobic contacts with the THIQ ring of LY3154207. It is replaced by a methionine in D5R and the bulky side chain of methionine may preclude LY3154207 and DETQ from binding to D5R due to steric clash (Fig. 1f). Consistent with the hypothesis, mutation of A139L in DRD1 largely decreased LY3154207 binding (Fig. 1h; Supplementary information, Figs. S7 and S9; Tables S3 and S4). Moreover, K138^4.40^ in TM4 forms a hydrogen bond with the long stretched tertiary alcohol of LY3154207, which is likely missing for DETQ due to the shorter linker between the tertiary alcohol and the THIQ ring in DETQ (Supplementary information, Fig. S10a). Consistently, LY3154207 showed a higher affinity to DRD1 than that of DETQ^6^.

We have also determined a cryo-EM structure of the DRD1-G_s_ complex with SKF82197 alone^8^. To investigate the mechanism of DRD1 allosteric modulation by LY3154207 and determine whether LY3154207 induces a different DRD1 conformation, we aligned the structures of DRD1 bound to both SKF81297 and LY3154207 and DRD1 bound to SKF81297 only. The overall DRD1 conformation in these two structures is highly similar (root mean square deviation 0.6 Å for Cα atoms) and the interactions between DRD1 and SKF81297 are nearly identical, but the binding poses of SKF81297 are slightly different and conformational differences can be observed in the allosteric binding pocket (Fig. 1e; Supplementary information, Fig. S10b). In the structure of DRD1 bound to LY3154207 and SKF81297, the binding pose of SKF81297 is 0.6 Å deeper compared to that of DRD1 bound to SKF81297, resulting in an extra hydrogen bond interaction between the *para*-hydroxyl group in the SKF81297 catechol ring and S198^5.42^ (Fig. 1e; Supplementary information, Fig. S10b), the wider spread polar interaction may trap SKF81297 in a more stable state than that with SKF81297 bound alone, thus stabilize the active state of the receptor. Furthermore, the extensive interactions between LY3154207 and ICL2 may stabilize the α-helical structure of ICL2, which is important for receptor activation and G_s_-coupling^12^. It is likely that LY3154207 potentiates the orthosteric agonist-induced DRD1 signaling by stabilizing the active state of DRD1, which is similar to the allosteric modulation of β_2_AR by Cmpd-6FA^11^.

In conclusion, the two structures reported here uncovered the unique binding modes of dopamine and the DRD1 potent PAM LY3154207, revealed the detailed mechanism of how DRD1 is occupied by both endogenous and synthetic agonists with different chemical scaffolds, and elucidated the potential allosteric regulation of DRD1 activity. Structure comparison demonstrated the important role of the DRD1 extended binding pocket in determining its ligand potency. In addition, the structure of LY3154207 bound to DRD1 presents a first view of the DRD1 PAM binding pocket. Together with mutagenesis results, our structures provide a framework for more efficient and subtype-selective ligand discovery targeting DRD1 for treating CNS diseases, both at the orthosteric ligand pocket and at the allosteric pocket.

## DATA AND MATERIALS AVAILABILITY

The cryo-EM density maps have been deposited in the Electron Microscopy Data Bank as EMD-23390 for DRD1-SKF81297-LY3154207-G_s_ and EMD-23391 for DRD1-dopamine-LY3154207-G_s_. The coordinates of the structures have been deposited in the Protein Data Bank as PDB 7LJC and PDB 7LJD for SKF81297-LY3154207 and dopamine-LY3154207 bound DRD1-G_s_ complexes, respectively.

## ACKNOWLEDGEMENTS

The cryo-EM datasets were collected at the Center of Cryo-Electron Microscopy, Zhejiang University and at the Center of Cryo-Electron Microscopy, Shanghai Institute of Materia Medica. This work was partially supported by the National Key R&D Programs of China 2018YFA0507002, Shanghai Municipal Science and Technology Major Project 2019SHZDZX02 and the Strategic Priority Research Program of Chinese Academy of Sciences XDB37030103 to H.E.X.; the National Natural Science Foundation of China 81922071, the National Key Basic Research Program of China 2019YFA0508800, Zhejiang Province Natural Science Fund for Excellent Young Scholars LR19H310001 and the Fundamental Research Funds for the Central Universities 2019XZZX001-01-06 to Y.Z.; Science and Technology Commission of Shanghai Municipal 20431900100 and Jack Ma Foundation 2020-CMKYGG-05 to H.J.; National Natural Science Foundation 31770796 and National Science and Technology Major Project 2018ZX09711002 to Y.J.; grants from the NIMH Psychoactive Drug Screening Program (X-P. H., L.Y., B.L.R.) and RO1MH112205 to B.K. and B.L.R.; NIH grant R35GM128641 to C.Z.

## AUTHOR CONTRIBUTIONS

Y.W.Z. and H.E.X initialed the project. Y.W.Z. designed the expression constructs, prepared protein samples for DRD1-G_s_ complexes for cryo-EM data collection, prepared the cryo-EM grids, performed data acquisition and structure determination of DRD1-SKF81297-LY3154207-G_s_, prepared the draft of manuscript and figures; B.K., X.P.H., and Y.L. performed cAMP assays and MiniG_s_ recruitment assays; H.B.Z. prepared the cryo-EM grids, conducted cryo-EM images collection and structure determination of DRD1-dopamine/ LY3154207-G_s_, participated in supplementary figures preparation; X.E.Z. refined and built the structure models; Y.W. assisted in preparation of DRD1 samples; X.C. performed ligand docking and simulation; Y.J. participated in project supervision; H.J. supervised X.C.; C.Z. participated in manuscript preparation; Y.W. synthesized the CID2886111 and LY3154207; B.L.R. supervised pharmacological and mutagenesis experiments and participated in manuscript writing; Y.Z. supervised H.B.Z. and the related EM works; H.E.X. conceived and supervised the project and wrote the manuscript with Y.W.Z., B.L.R. and C.Z.

---

**Fig. S1.**
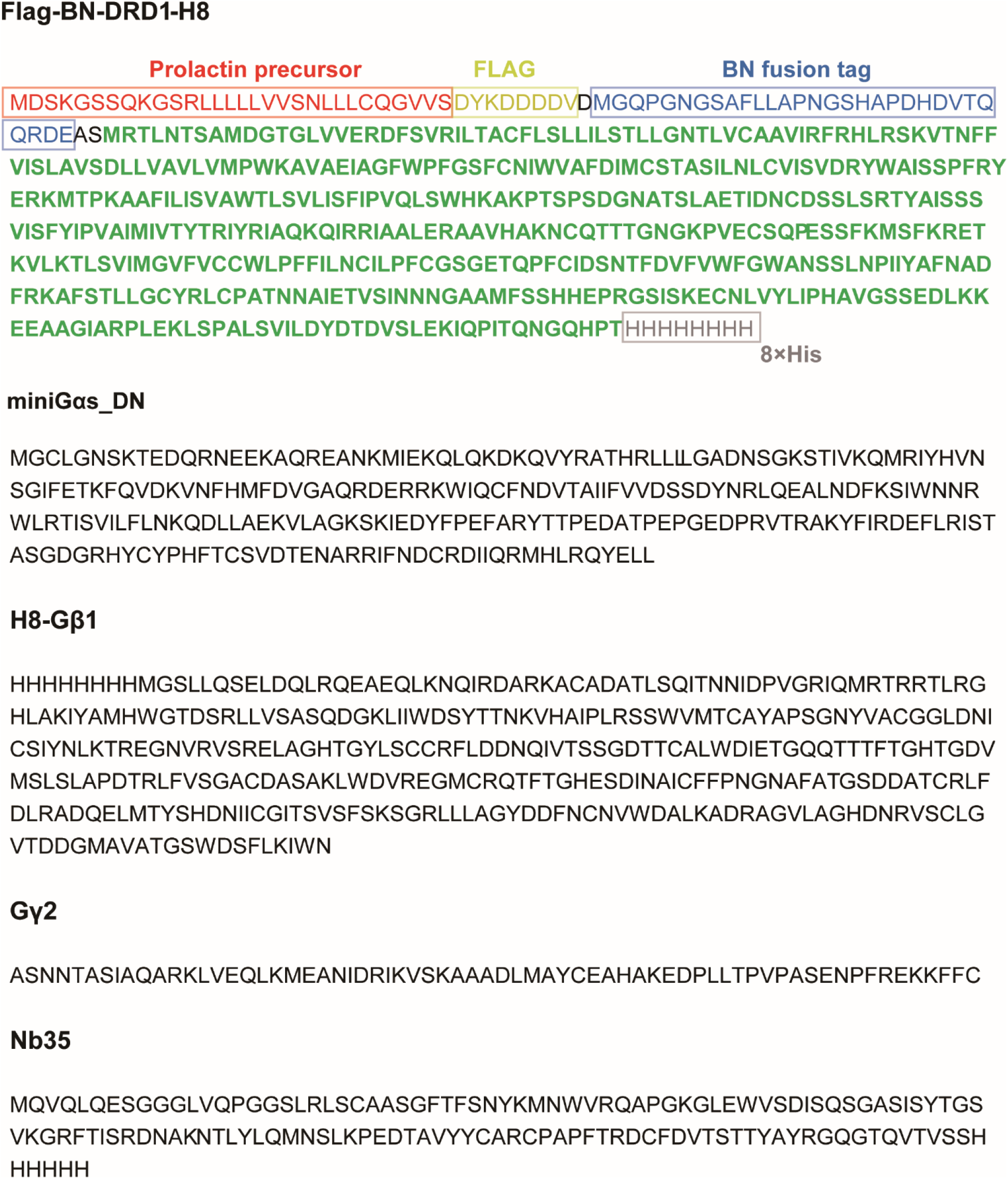
The amino acid sequences of DRD1-G_s_ complex components used in this study. The DRD1 sequence is showed in green.

**Fig. S2.**
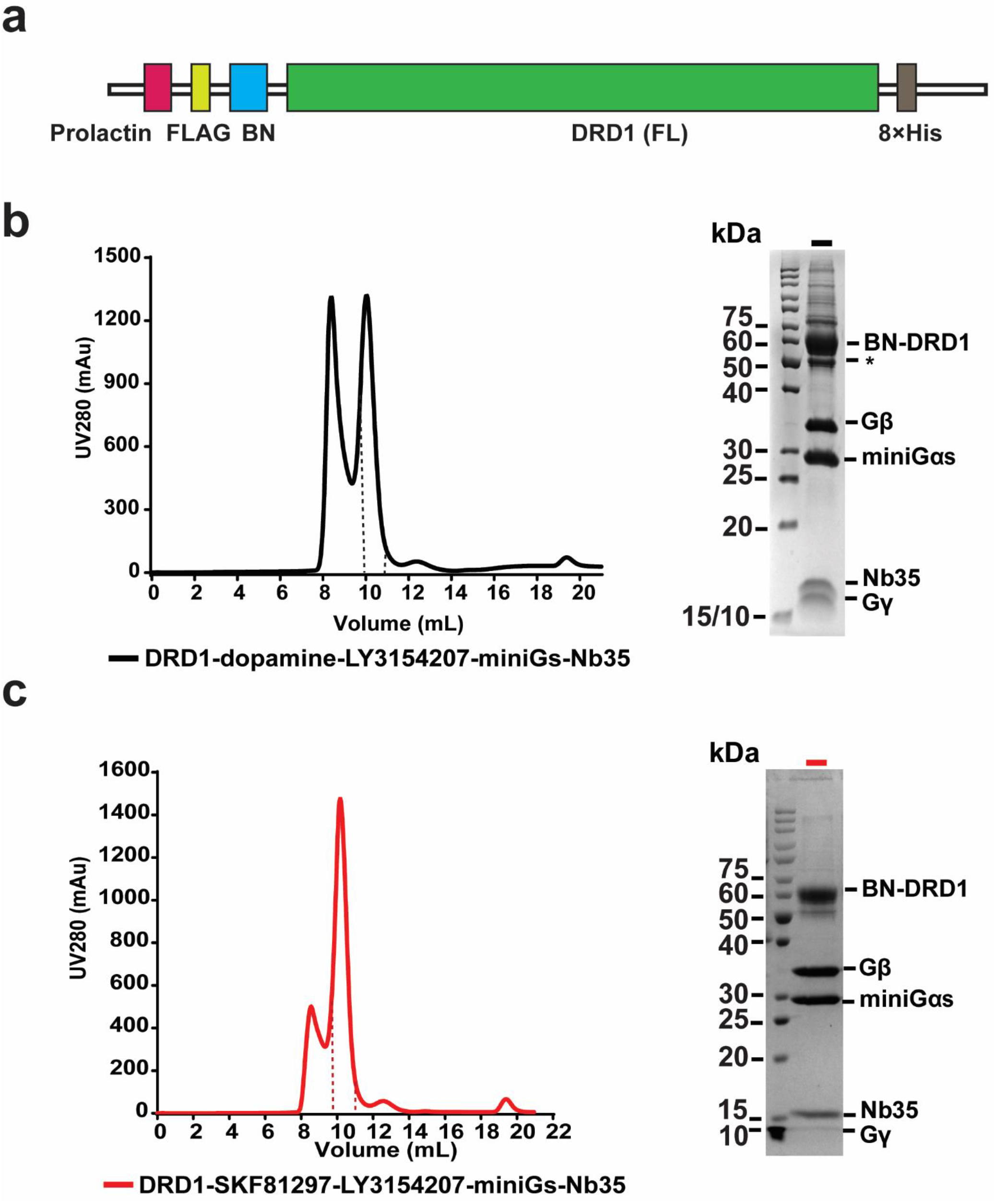
Construct and preparation of DRD1-miniG_s_ cryo-EM samples. **a** A cartoon model of the DRD1 construct used for cryo-EM studies. BN: β_2_AR N-terminal tail region; Prolactin: prolactin signal peptide sequence. **b** Size exclusion chromatography profile and SDS-PAGE analysis of DRD1-dopamine/ LY3154207-miniG_s_ complex. Fractions between the two dashed line were collected and concentrated for further cryo-EM grids preparation. **c** Size exclusion chromatography separation and SDS-PAGE analysis of the DRD1-SKF81297/LY3154207-miniG_s_ complex. Protein fractions between the two dashed line were collected and concentrated for subsequent cryo-EM studies.

**Fig. S3.**
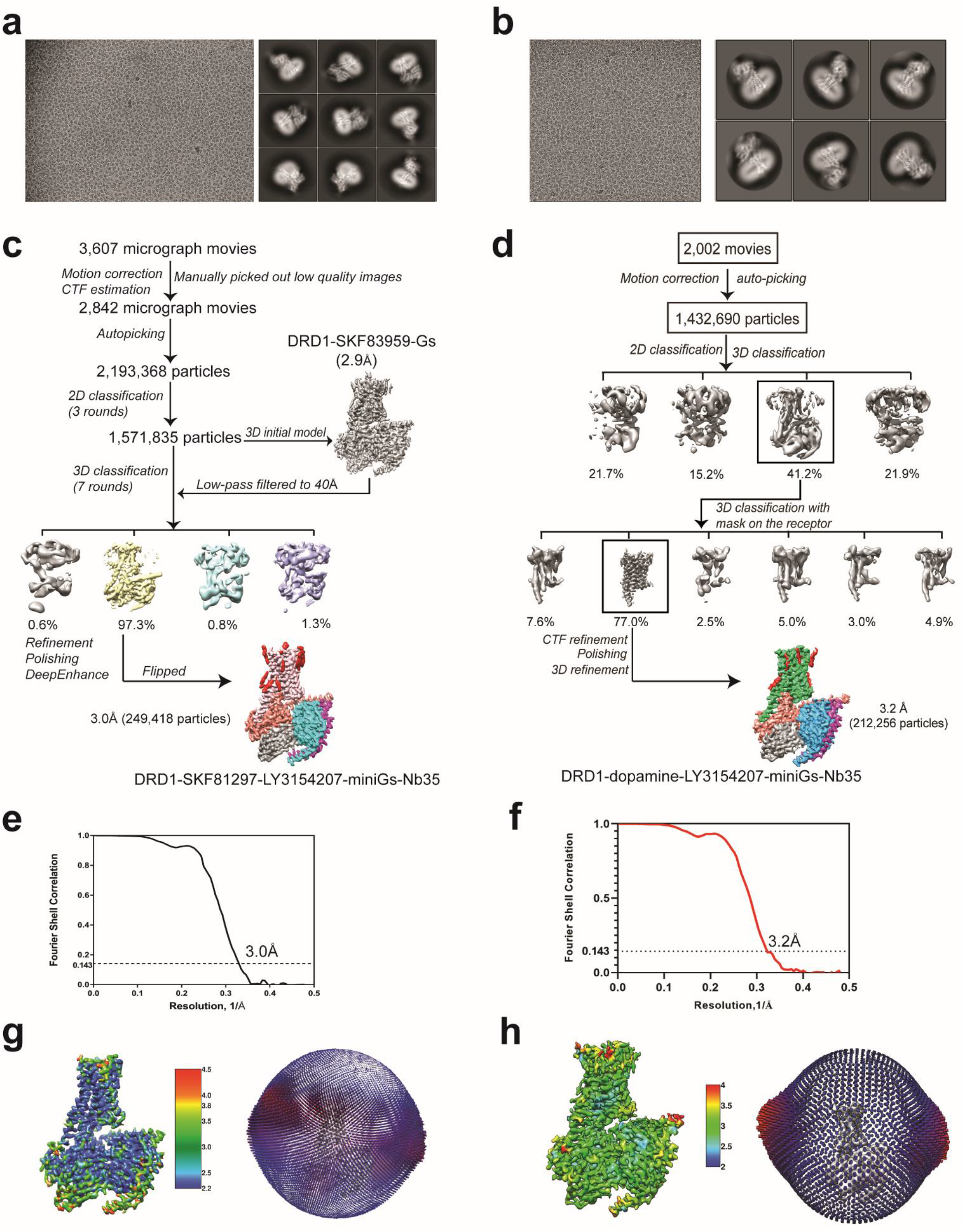
Single particle cryo-EM workflows for structure determination of DRD1-miniG_s_ complexes. **a - b** Representative cryo-EM micrographs and 2D classification averages of DRD1-SKF81297/ LY3154207-miniG_s_ **(a)** and DRD1-dopamine/ LY3154207-miniG_s_ **(b)**. The 2D averages show divergent secondary structural features from different views. **c-f** Cryo-EM data processing flowcharts of DRD1-SKF81297/ LY3154207-miniG_s_ **(c)** and DRD1-dopamine/ LY3154207-miniG_s_ **(d)** by Relion 3.0, including the final processed global density maps and the ‘Gold-standard’ Fourier shell correlation (FSC) curves of DRD1-SKF81297/ LY3154207-miniG_s_ **(e)** and DRD1-dopamine/ LY3154207-miniG_s_ **(f)**, respectively. The global resolution defined at the FSC=0.143 is 3.0 Å for DRD1-SKF81297/ LY3154207-miniG_s_ and 3.2 Å for DRD1-dopamine/ LY3154207-miniG_s_. **g - h** The global density maps of DRD1-SKF81297/ LY3154207-miniG_s_ **(g)** and DRD1-dopamine/ LY3154207-miniG_s_ **(h)** colored by local resolution (Å), along with the angle distribution maps of particle orientations. The density maps are shown at thresholds of 0.083 and 0.045 for SKF81297/ LY3154207 and dopamine/ LY3154207-bound complexes, respectively.

**Fig. S4.**
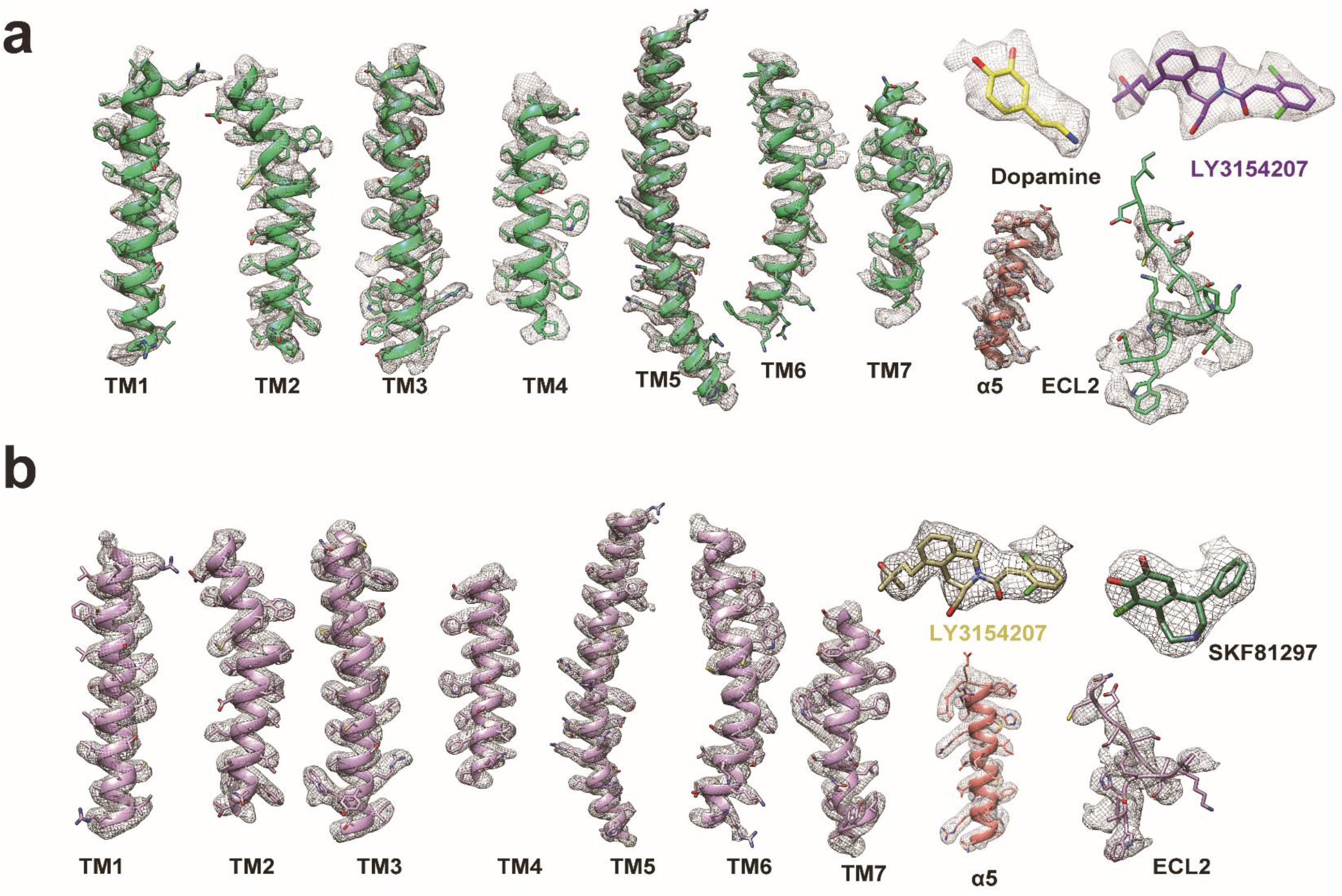
Local cryo-EM density maps of DRD1-miniG_s_ complexes. Cryo-EM density maps of TM1-TM7, ligands and ECL2 of DRD1 and α5 helix of miniG_αs_ in the DRD1-dopamine/LY3154207-miniG_s_ structure **(a)** and the DRD1-SKF81297/ LY3154207-miniG_s_ structure **(b)**.

**Fig. S5.**
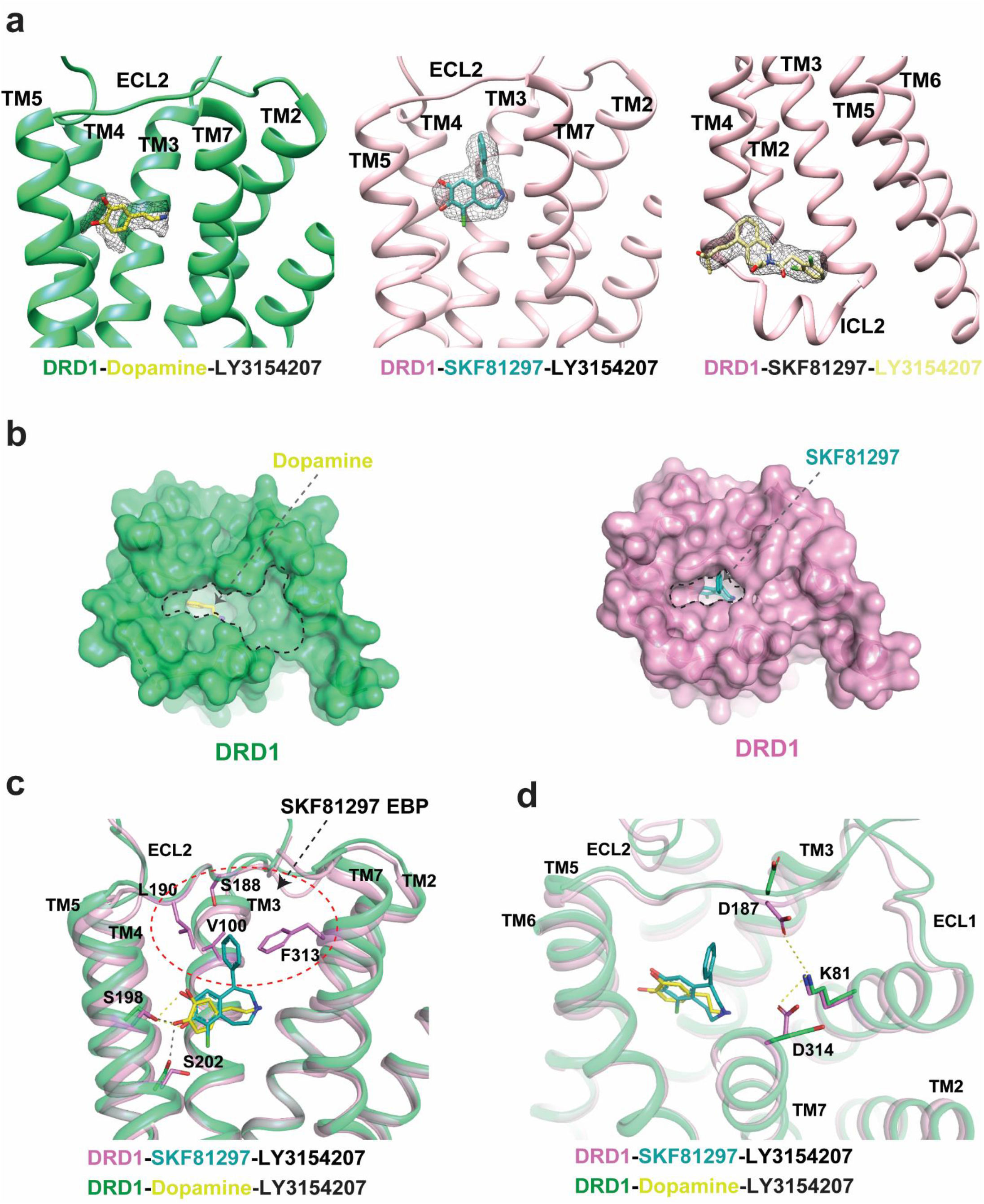
Topologies of ligand binding pockets of DRD1-miniG_s_ complexes. **a** The ligand binding poses of dopamine in the DRD1-dopamine/ LY3154207-miniGs structure and of SKF81297 and LY3154207 in the DRD1-SKF81297/ LY3154207-miniG_s_ structure. The density of each ligand was clear and shown as mesh. **b** Topologies of extracellular vestibules of DRD1 activated by dopamine and SKF81297, respectively. The dopamine-bound DRD1 adopts a more open extracellular vestibules compared to the SKF81297-bound DRD1. **c** Comparison of interactions and binding poses in the DRD1 ligand binding pockets of dopamine and SKF81297. The SKF81297 compound forms extended interactions with the EBP of DRD1, which is mainly formed by residues from TM3, ECL2 and TM7. The EBP region is circled with a red dashed line. Hydrogen bonds are shown as black dashed lines in the dopamine-bound DRD1 structure; in SKF81297-bound DRD1, the hydrogen bond interactions are shown as yellow dashed line. **d** Comparison of conformational arrangements between residues K81, D187 and D314 in the DRD1-dopamine structure and the DRD1-SKF81297/ LY3154207 structure. D187 points toward K81 and forms a polar interaction network with K81 and D314 in the DRD1-SKF81297/ LY3154207 structure, but not in the DRD1-dopamine/ LY3154207 structure. The polar interactions are shown as yellow dashed lines.

**Fig. S6.**
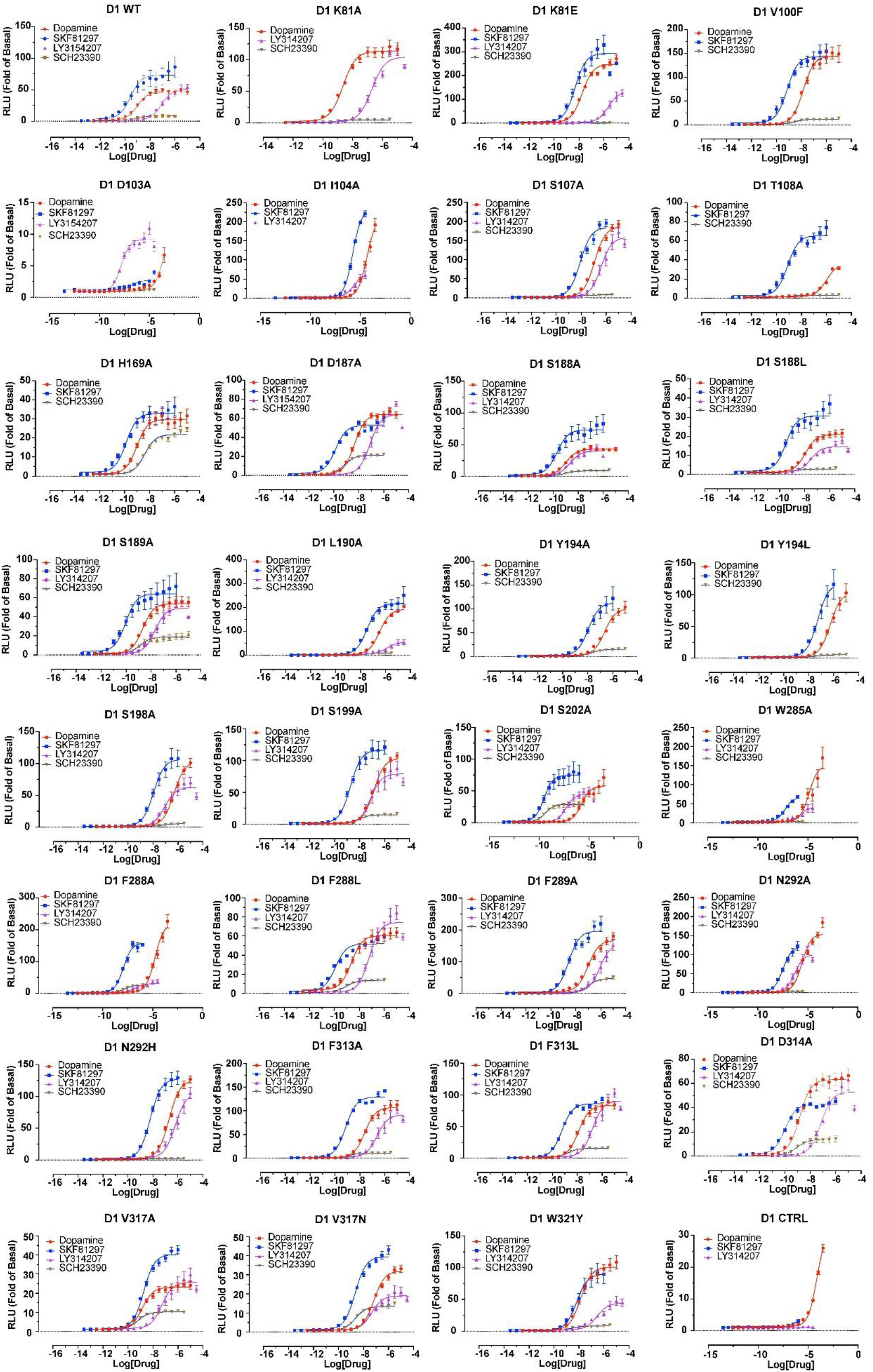
Comparison of dopamine, SKF81297, and LY3154207 in cAMP accumulation assays with WT DRD1 and DRD1 mutants. Normalized dose response curves of cAMP accumulation assay (Glosensor) data of WT DRD1 and DRD1 mutants activated by dopamine, SKF81297 and LY3154207, respectively. Data are presented as mean values ± SEM with a minimum of two technical replicates and N = 3 biological replicates.

**Fig. S7.**
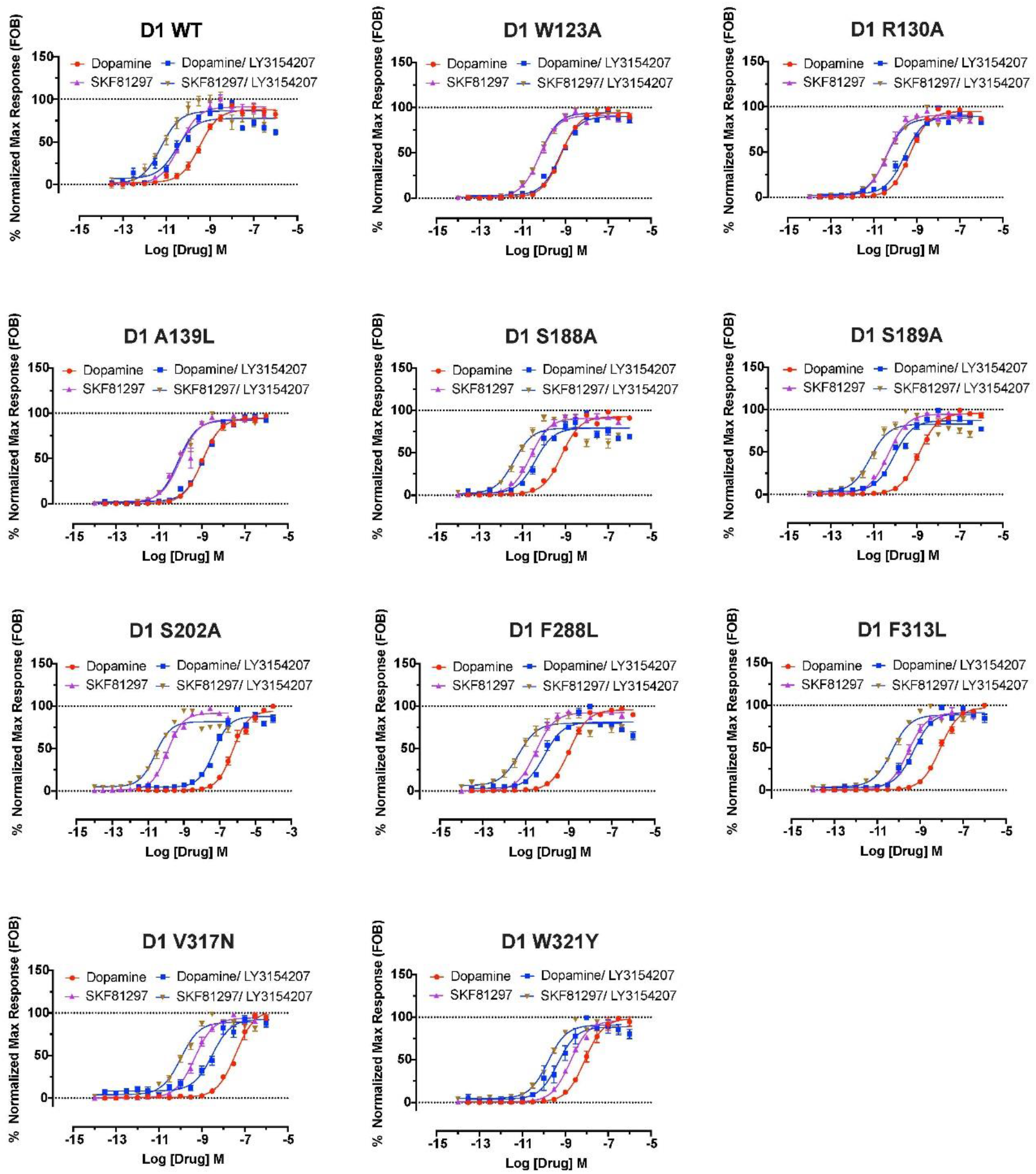
Comparison of dopamine and SKF81297 cAMP accumulation assays to WT DRD1 and DRD1 mutants in the presence or absence of 30 nM LY3154207. Data are presented as mean values ± SEM with a minimum of two technical replicates and N = 3 biological replicates. F-test analysis of pEC50 values of the following mutants were not statistically different when comparing dose response curves in the presence and absence of 30 nM LY3154207 indicative of diminished allosteric effect at these mutants. F-test analysis (p < 0.05), F_(1,258)_W123A^Dopamine^ = 1.263, p = 0.2622; F_(1,168)_W123A^SKF81297^ = 0.1890, p = 0.6643; F_(1,279)_R130A^SKF81297^ = 0.02710, p = 0.8694; F_(1,258)_A139L^Dopamine^ = 0.5500, p = 0.4591; F_(1,279)_A139L^SKF81297^ = 1.525, p = 0.2180. WT F-test analysis for comparison which was statistically different, F_(1,362)_WT^Dopamine^ = 113.5, p < 0.0001 and F_(1,354)_WT^SKF81297^ = 36.11, p < 0.0001. See Table S3 for additional statistical information. Data related to Fig. 1g and 1h.

**Fig. S8.**
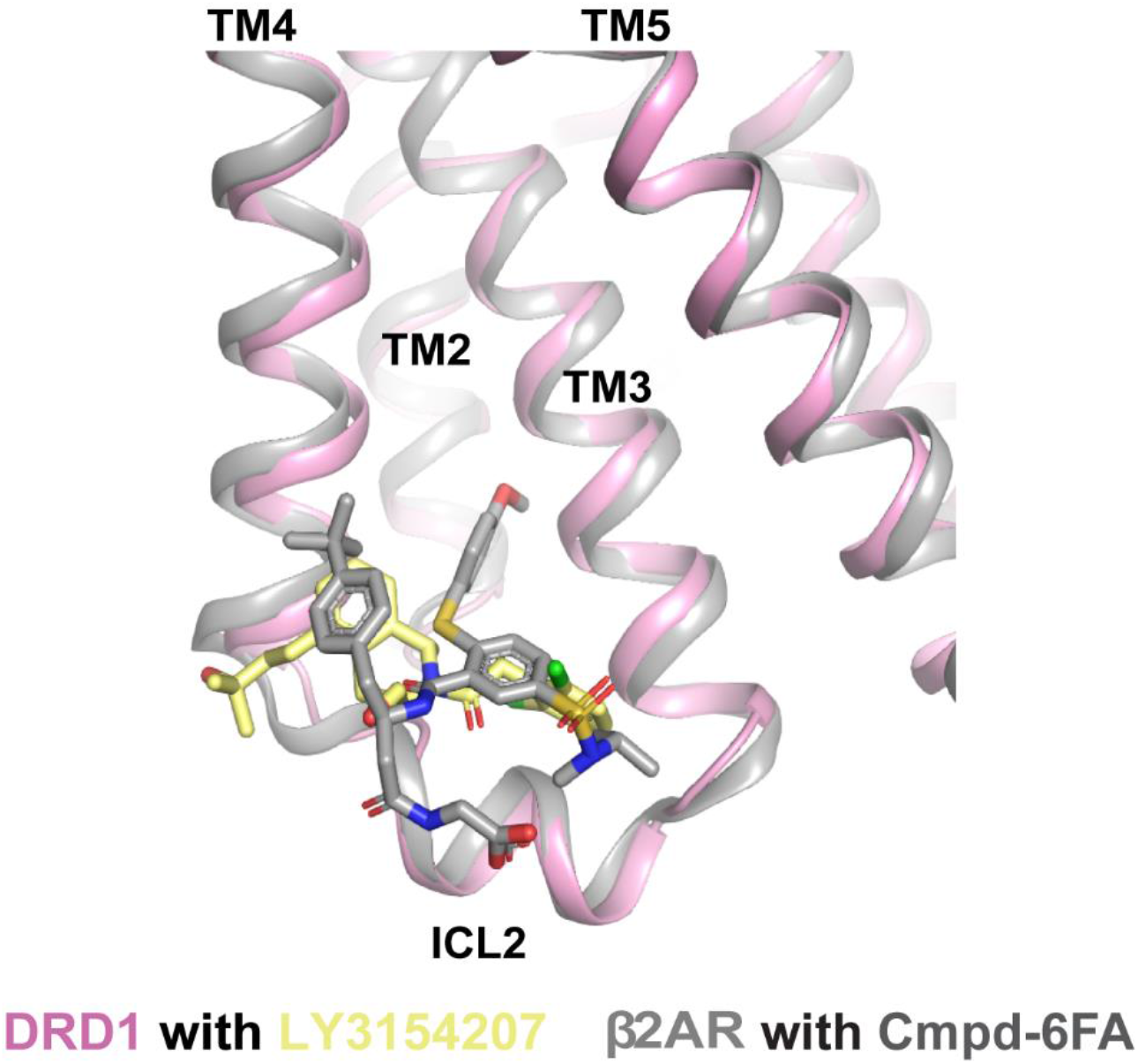
Structural alignment of the PAM-binding sites in DRD1 and β_2_AR. The structure of β_2_AR bound with Cmpd-6FA (PDB code: 6N48) is colored gray, DRD1 is colored pink and the DRD1 PAM compound LY3154207 is colored light yellow.

**Fig. S9.**
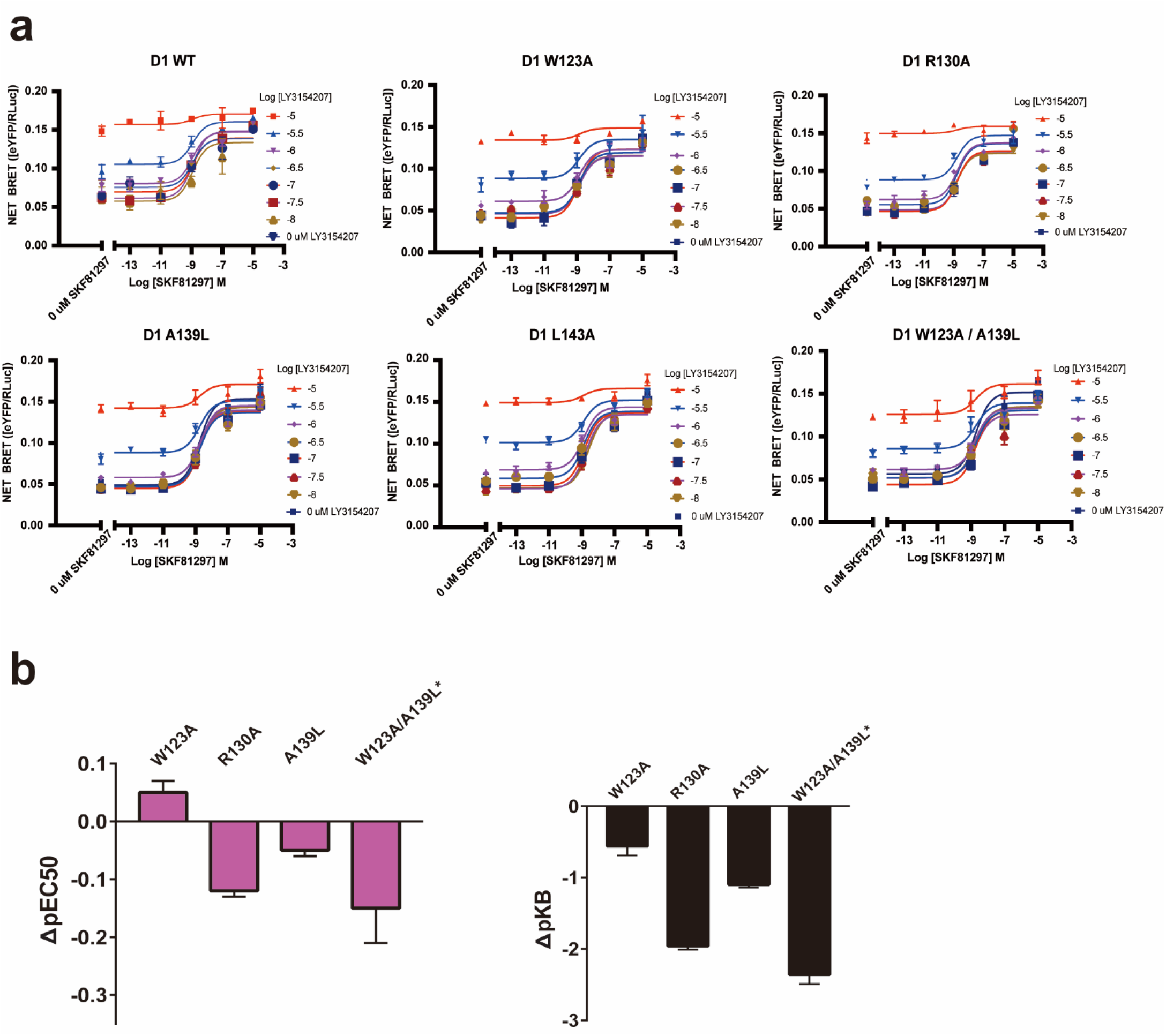
Mutations of D1R allosteric site residues affect LY3154207 allosteric properties. BRET Venus-MiniG_s_ recruitment SKF81297 and LY3154207 dose response curves **(a)** and bar charts **(b)** for identified D1R WT and mutations. F-test analysis (p < 0.05) comparing 0 μM and 30 nM (log_10_ = −7.5) LY3154207 show only WT is statistically different indicating an allosteric effect for WT and diminished allosteric effect for identified allosteric site mutants. F-test results - F_(3,63)_WT = 5.798, p = 0.0014; F_(3,63)_W123A = 1.380, p = 0.2572; F_(3,63)_R130A = 0.4484, p = 0.7193; F_(3,63)_A139L = 0.8458, p = 0.4740; F_(3,63)_L143A = 1.814, p = 0.1536; F_(3,38)_W123A/A139L = 1.873, p = 0.1507.

**Fig. S10.**
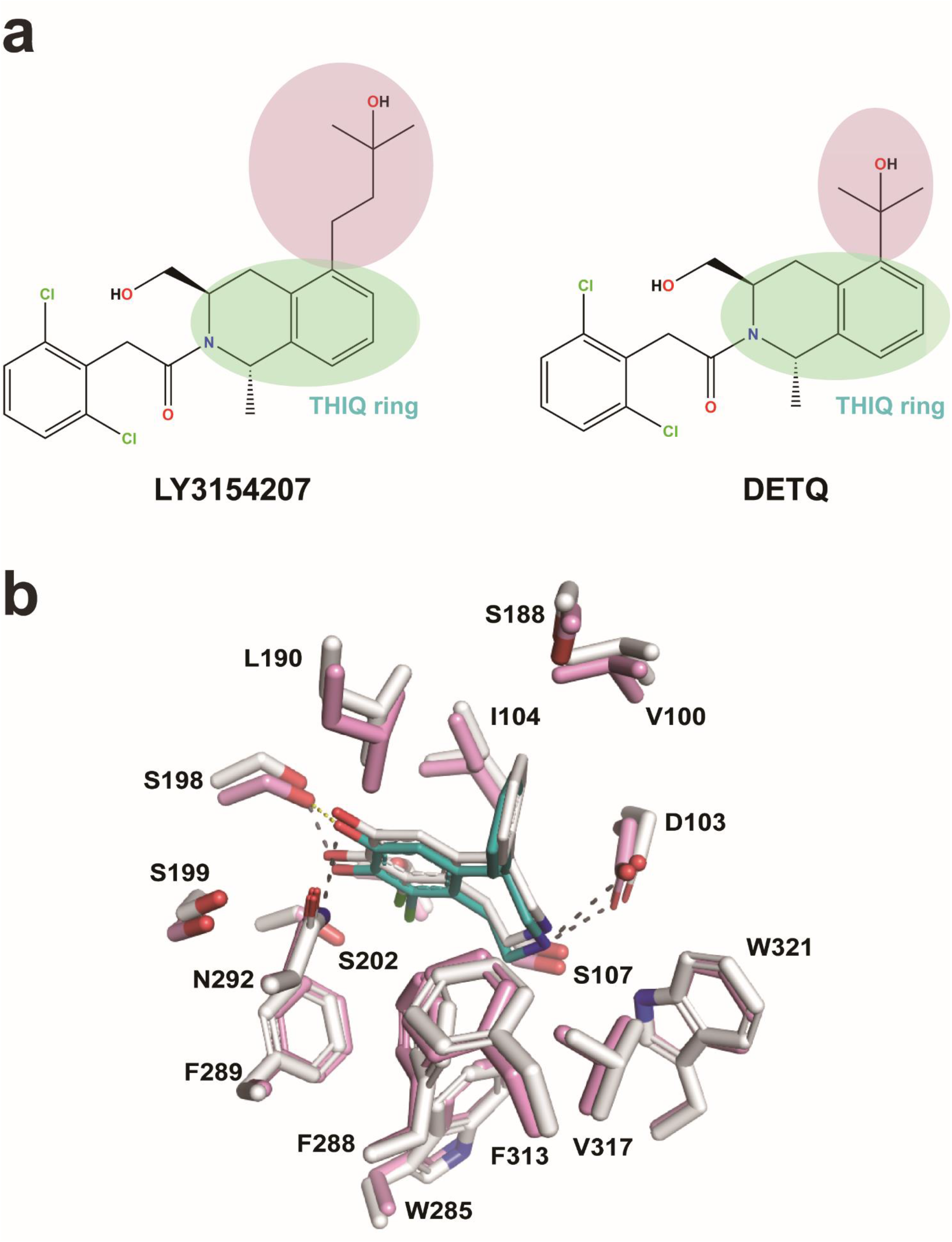
Superposition of the orthosteric pocket in active DRD1 in presence or absence of LY3154207. **a** Chemical structures of DRD1 PAM compounds DETQ and LY3154207. The THIQ ring is marked with a light green circle while the alkyl linker between the C5 tertiary alcohol and THIQ ring and C5 tertiary alcohol is marked with a light pink circle in both structures. The alkyl linker in LY3154207 is longer than that of DETQ, which is the only difference between these two DRD1 PAMs. **b** Preserved SKF81297 orthosteric binding pocket in active DRD1 with or without LY3154207. The residues participating in interaction with agonist are well overlapped while the binding pose of SKF81297 in LY3154207-bound active DRD1, which is deeper than that of active DRD1 alone, leading to a closer polar interaction with S202. The DRD1-SKF81297 structure without PAM is colored white, the DRD1 and SKF81297 in the DRD1-SKF81297/ LY3154207 structure ware colored pink and steal, respectively. The shared hydrogen bond interactions are shown as black dashed line while the additional hydrogen interaction between S198^5.42^ and SKF81297 in the DRD1-SKF81297/ LY3154207 structure is shown as yellow dashed line.

**Table S1.**
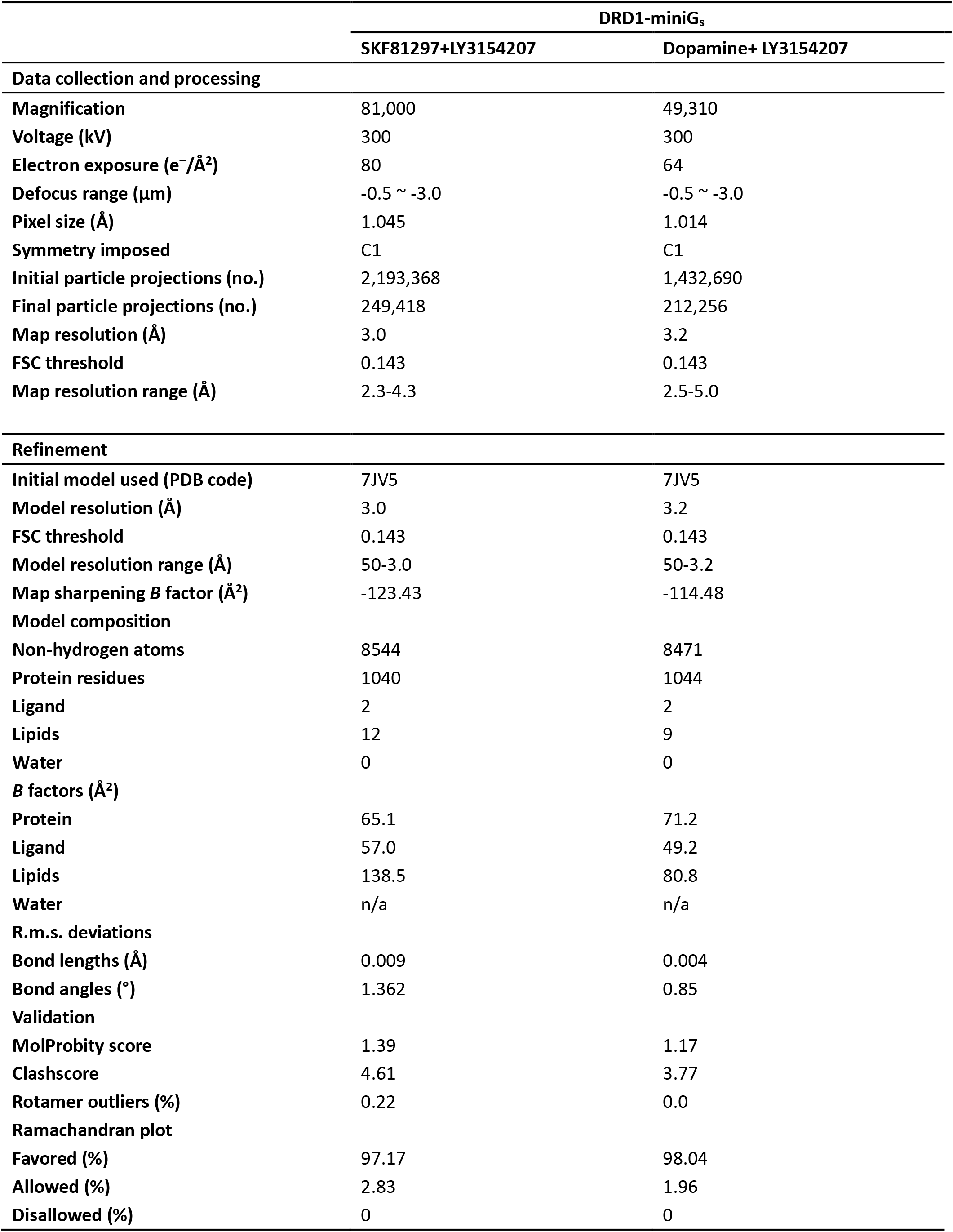
Cryo-EM data collection, refinement and validation statistics.

**Table S2.** cAMP accumulation results of WT DRD1 and DRD1 mutants in orthosteric site. Data are presented as mean values ± SEM with a minimum of two technical replicates and N = 3 biological replicates. Greek letter delta (Δ) for the difference (ΔpEC50) and Fold of Basal (ΔFOB) when compared with the wild-type receptor values. ND – Not Determined. Data related to Fig. S6.

| D1R | Dopamine |  |  |  | SKF81297 |  |  |  | LY3154207 |  |  |  | SCH23390 |  |  |  |
| --- | --- | --- | --- | --- | --- | --- | --- | --- | --- | --- | --- | --- | --- | --- | --- | --- |
|  | Fold of Basal (FOB) | ΔFOB | pEC50 | ΔpEC50 | Fold of Basal (FOB) | ΔFOB | pEC50 | ΔpEC50 | Fold of Basal (FOB) | ΔFOB | pEC50 | ΔpEC50 | Fold of Basal (FOB) | ΔFOB | pEC50 | ΔpEC50 |
| WT | 51.93 ± 7.01 | 0.00 | 9.12 ± 0.16 | 0.00 | 74.57 ± 30.15 | 0.00 | 9.85 ± 0.28 | 0.00 | 53.34 ± 13.1 | 0.00 | 7.02 ± 0.28 | 0.00 | 9.47 ± 1.24 | 0.00 | 9.04 ± 0.13 | 0.00 |
| K81A | 118.73 ± 19.08 | 66.80 | 8.72 ± 0.23 | -0.40 | ND | ND | ND | ND | 107.97 ± 10.05 | 54.63 | 6.7 ± 0.4 | -0.32 | 5.45 ± 0.24 | -4.01 | 8.85 ± 0.08 | -0.20 |
| K81E | 262.73 ± 28.82 | 210.80 | 7.69 ± 0.18 | -1.43 | 286.33 ± 97.96 | 211.76 | 8.35 ± 0.28 | -1.50 | 192.81 ± 41.47 | 139.47 | 5.09 ± 0.59 | -1.94 | 2.7 ± 0.54 | -6.77 | 8.17 ± 0.03 | -0.87 |
| V100F | 192.7 ± 69.49 | 140.77 | 8.09 ± 0.21 | -1.03 | 137.85 ± 39.77 | 63.28 | 9.07 ± 0.22 | -0.78 | ND | ND | ND | ND | 9.98 ± 1.02 | 0.51 | 8.61 ± 0.14 | -0.43 |
| D103A | 15.82 ± 5.73 | -36.11 | 3.32 ± 0.04 | -5.81 | 7.05 ± 3.4 | -67.52 | 4.77 ± 1.13 | -5.08 | 9.54 ± 1.83 | -43.80 | 7.73 ± 0.21 | 0.71 | ND | ND | ND | ND |
| I104A | 317.7 ± 53.92 | 265.77 | 4.13 ± 0.19 | -4.99 | 297.19 ± 57.92 | 222.62 | 5.86 ± 0.14 | -3.99 | 83.09 ± 60.02 | 29.75 | 6.23 ± 0.92 | -0.79 | ND | ND | ND | ND |
| S107A | 200.52 ± 24.88 | 148.59 | 6.89 ± 0.16 | -2.23 | 188.29 ± 44.68 | 113.73 | 8.27 ± 0.25 | -1.58 | 167.08 ± 26.73 | 113.74 | 6.23 ± 0.28 | -0.79 | 7.64 ± 0.62 | -1.82 | 8.61 ± 0.06 | -0.44 |
| T108A | 29.7 ± 9.4 | -22.23 | 5.85 ± 0.19 | -3.27 | 68.57 ± 17.33 | -6.00 | 8.76 ± 0.19 | -1.09 | ND | ND | ND | ND | 3.43 ± 0.52 | -6.03 | 8.91 ± 0.12 | -0.13 |
| H169A | 30.03 ± 7.91 | -21.90 | 9.11 ± 0.24 | -0.01 | 34.17 ± 9.77 | -40.40 | 10.05 ± 0.32 | 0.20 | ND | ND | ND | ND | 21.33 ± 2.3 | 11.86 | 8.3 ± 0.07 | -0.74 |
| D187A | 65.11 ± 6.61 | 13.18 | 8.35 ± 0.13 | -0.77 | ND | ND | ND | ND | 66.84 ± 3.56 | 13.50 | 6.94 ± 0.32 | -0.08 | 21.93 ± 2.11 | 12.46 | 8.96 ± 0.12 | -0.09 |
| S188A | 43.46 ± 4.6 | -8.47 | 9.05 ± 0.16 | -0.07 | 54.34 ± 21.6 | -20.22 | 10.17 ± 0.26 | 0.32 | 39.81 ± 5.12 | -13.53 | 7.03 ± 0.22 | 0.01 | 9.18 ± 1.22 | -0.29 | 8.92 ± 0.16 | -0.12 |
| S188L | 21.95 ± 3.81 | -29.98 | 7.98 ± 0.2 | -1.14 | 25.63 ± 5.63 | -48.94 | 9.76 ± 0.4 | -0.09 | 15.69 ± 3.29 | -37.65 | 7.44 ± 0.22 | 0.42 | 3.17 ± 0.18 | -6.30 | 8.32 ± 0.37 | -0.73 |
| S189A | 59.72 ± 13.41 | 7.79 | 8.66 ± 0.11 | -0.46 | 70.18 ± 29.28 | -4.39 | 10.14 ± 0.32 | 0.29 | 51.75 ± 2.51 | -1.59 | 7.05 ± 0.47 | 0.03 | 19.86 ± 6.77 | 10.40 | 8.92 ± 0.14 | -0.12 |
| L190A | 258.06 ± 31.62 | 206.13 | 6.39 ± 0.49 | -2.73 | 220.89 ± 40.72 | 146.32 | 7.47 ± 0.08 | -2.38 | 89.88 ± 34.83 | 36.54 | 5.5 ± 1.08 | -1.52 | 10.43 ± 0.88 | 0.96 | 7.34 ± 0.14 | -1.70 |
| Y194A | 113.57 ± 18.62 | 61.64 | 6.6 ± 0.15 | -2.53 | 117.01 ± 43.76 | 42.44 | 8.11 ± 0.33 | -1.74 | ND | ND | ND | ND | 12.72 ± 0.96 | 3.25 | 7.56 ± 0.05 | -1.48 |
| Y194L | 124.91 ± 45.48 | 72.98 | 6.32 ± 0.33 | -2.80 | 133.99 ± 70.23 | 59.42 | 7.38 ± 0.28 | -2.47 | ND | ND | ND | ND | 4.91 ± 0.67 | -4.56 | 7.41 ± 0.23 | -1.63 |
| S198A | 94.25 ± 23.63 | 42.32 | 6.09 ± 0.31 | -3.03 | 98.06 ± 40.31 | 23.49 | 8.11 ± 0.19 | -1.74 | 63.98 ± 16.46 | 10.64 | 6.99 ± 0.25 | -0.03 | 5.97 ± 0.79 | -3.50 | 7.07 ± 0.1 | -1.97 |
| S199A | 107.63 ± 6.91 | 55.69 | 6.86 ± 0.07 | -2.26 | 116.04 ± 28.56 | 41.47 | 8.76 ± 0.15 | -1.09 | 84.11 ± 23.54 | 30.77 | 6.78 ± 0.54 | -0.25 | 16.06 ± 2.47 | 6.59 | 8.01 ± 0.17 | -1.04 |
| S202A | 67.82 ± 15.78 | 15.89 | 5.46 ± 0.32 | -3.66 | 73.94 ± 32.29 | -0.63 | 9.59 ± 0.18 | -0.26 | 48.37 ± 7.65 | -4.97 | 7.09 ± 0.23 | 0.07 | 29.25 ± 7.07 | 19.78 | 9.39 ± 0.12 | 0.34 |
| W285A | 368.54 ± 143.29 | 316.61 | 4.43 ± 0.33 | -4.69 | ND | ND | ND | ND | 72.48 ± 59.8 | 19.14 | 6.25 ± 0.88 | -0.77 | 3.6 ± 0.19 | -5.87 | 8.8 ± 0.1 | -0.24 |
| F288A | 199.38 ± 47.89 | 147.45 | 4.56 ± 0.31 | -4.56 | 101.59 ± 33.5 | 27.03 | 8.09 ± 0.15 | -1.76 | 67.38 ± 46.64 | 14.04 | 6.19 ± 1 | -0.83 | 27.75 ± 6.87 | 18.28 | 7.81 ± 0.12 | -1.24 |
| F288L | 66.52 ± 11.33 | 14.59 | 8.9 ± 0.43 | -0.22 | 63.1 ± 22.14 | -11.47 | 9.58 ± 0.9 | -0.27 | 86.61 ± 15.05 | 33.27 | 6.92 ± 0.43 | -0.10 | 13.52 ± 4.38 | 4.05 | 8.88 ± 0.04 | -0.16 |
| F289A | 174.41 ± 32.53 | 122.48 | 7.25 ± 0.4 | -1.87 | ND | ND | ND | ND | 161.09 ± 19.76 | 107.74 | 5.86 ± 0.44 | -1.16 | 51.13 ± 11.74 | 41.67 | 7.43 ± 0.19 | -1.62 |
| N292A | 222.7 ± 39.57 | 170.77 | 4.73 ± 0.27 | -4.39 | 118.3 ± 29.95 | 43.74 | 7.57 ± 0.19 | -2.28 | 112.15 ± 10.71 | 58.81 | 6.01 ± 0.45 | -1.01 | 2.45 ± 0.14 | -7.01 | 8.28 ± 0.32 | -0.76 |
| N292H | 120.63 ± 18.62 | 68.70 | 6.78 ± 0.14 | -2.34 | 108.49 ± 46.96 | 33.92 | 8.35 ± 0.12 | -1.50 | 110.31 ± 0.61 | 56.97 | 6 ± 0.5 | -1.02 | 4.6 ± 1.87 | -4.86 | 6.64 ± 1.52 | -2.40 |
| F313A | 111.41 ± 21.82 | 59.48 | 7.62 ± 0.13 | -1.50 | ND | ND | ND | ND | 104.85 ± 15.84 | 51.51 | 6.24 ± 0.44 | -0.78 | 11.04 ± 1.64 | 1.57 | 8.83 ± 0.05 | -0.22 |
| F313L | 85.37 ± 13.06 | 33.44 | 7.89 ± 0.1 | -1.23 | 79.86 ± 13.51 | 5.30 | 9.29 ± 0.12 | -0.56 | 102.7 ± 3.54 | 49.36 | 6.44 ± 0.39 | -0.58 | 16.76 ± 2.93 | 7.29 | 8.72 ± 0.05 | -0.32 |
| D314A | 56.68 ± 5.97 | 4.75 | 7.38 ± 0.09 | -1.75 | ND | ND | ND | ND | 61.34 ± 19.33 | 8.00 | 7 ± 0.29 | -0.02 | 7.4 ± 1.24 | -2.07 | 9.14 ± 0.09 | 0.10 |
| V317A | 23.8 ± 4.06 | -28.13 | 8.89 ± 0.12 | -0.24 | 41.76 ± 4.14 | -32.80 | 8.7 ± 0.03 | -1.16 | 27.76 ± 7.1 | -25.58 | 7.08 ± 0.28 | 0.06 | 10.39 ± 0.49 | 0.93 | 9.17 ± 0.06 | 0.13 |
| V317N | 34.13 ± 4.23 | -17.80 | 7.11 ± 0.15 | -2.01 | 35.39 ± 4.01 | -39.18 | 8.72 ± 0.1 | -1.13 | 19.77 ± 6.35 | -33.57 | 7.37 ± 0.19 | 0.34 | 14.68 ± 0.85 | 5.21 | 8.47 ± 0.06 | -0.57 |
| W321Y | 105.53 ± 22.86 | 53.60 | 7.86 ± 0.2 | -1.26 | 118.44 ± 32.63 | 43.87 | 8.48 ± 0.18 | -1.37 | 51.3 ± 15.73 | -2.04 | 6.37 ± 0.04 | -0.66 | 7.54 ± 1.57 | -1.93 | 8.14 ± 0.15 | -0.90 |

**Table S3.**
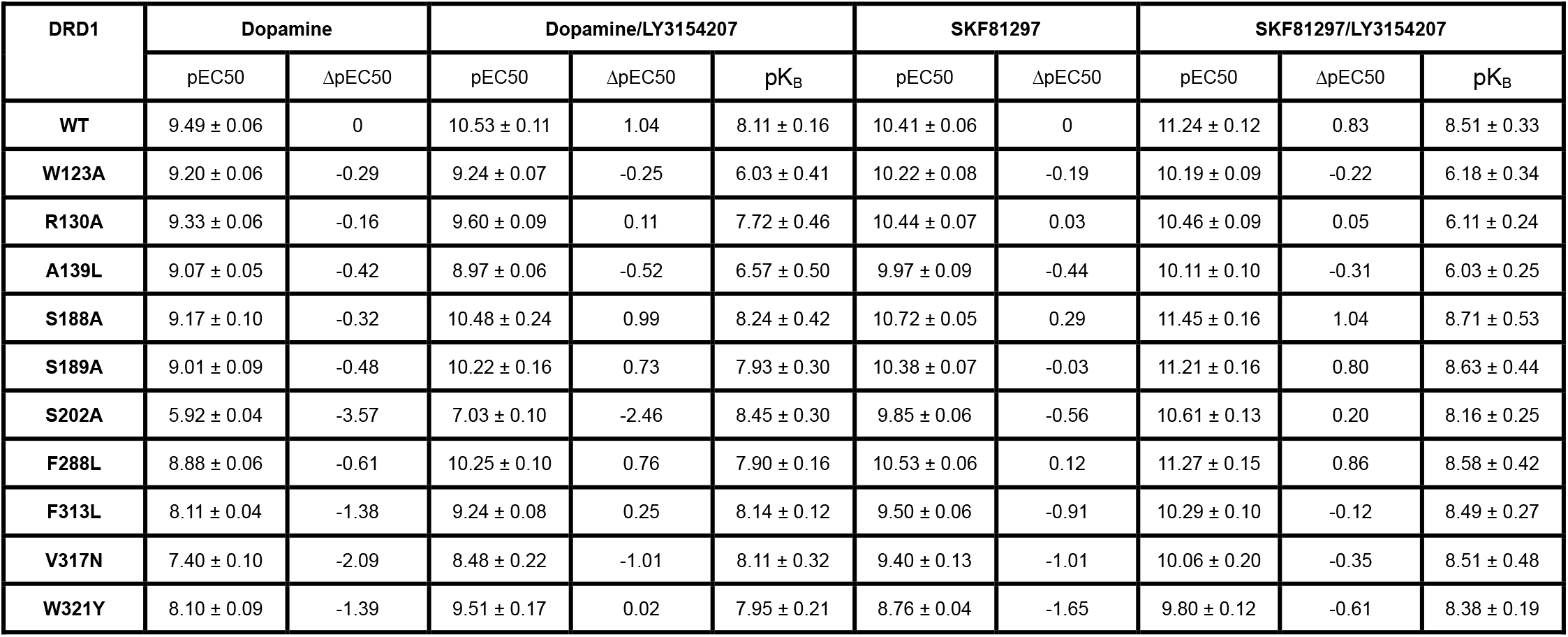
G_S_-mediated cAMP accumulation analysis of WT DRD1 and DRD1 mutants with or without PAM LY3154207. Data are presented as mean values ± SEM with a minimum of two technical replicates and N = 3 biological replicates. Greek letter delta (Δ) for the difference (ΔpEC50) when compared with the wild-type receptor values in the absence of 30 nM LY3154207. Data related to Fig. 1g and 1h.

| DRD1 | Dopamine |  | Dopamine/LY3154207 |  |  | SKF81297 |  | SKF81297/LY3154207 |  |  |
| --- | --- | --- | --- | --- | --- | --- | --- | --- | --- | --- |
|  | pEC<sub>50</sub> | ΔpEC<sub>50</sub> | pEC<sub>50</sub> | ΔpEC<sub>50</sub> | pK<sub>B</sub> | pEC<sub>50</sub> | ΔpEC<sub>50</sub> | pEC<sub>50</sub> | ΔpEC<sub>50</sub> | pK<sub>B</sub> |
| WT | 9.49 ± 0.06 | 0 | 10.53 ± 0.11 | 1.04 | 8.11 ± 0.16 | 10.41 ± 0.06 | 0 | 11.24 ± 0.12 | 0.83 | 8.51 ± 0.33 |
| W123A | 9.20 ± 0.06 | -0.29 | 9.24 ± 0.07 | -0.25 | 6.03 ± 0.41 | 10.22 ± 0.08 | -0.19 | 10.19 ± 0.09 | -0.22 | 6.18 ± 0.34 |
| R130A | 9.33 ± 0.06 | -0.16 | 9.60 ± 0.09 | 0.11 | 7.72 ± 0.46 | 10.44 ± 0.07 | 0.03 | 10.46 ± 0.09 | 0.05 | 6.11 ± 0.24 |
| A139L | 9.07 ± 0.05 | -0.42 | 8.97 ± 0.06 | -0.52 | 6.57 ± 0.50 | 9.97 ± 0.09 | -0.44 | 10.11 ± 0.10 | -0.31 | 6.03 ± 0.25 |
| S188A | 9.17 ± 0.10 | -0.32 | 10.48 ± 0.24 | 0.99 | 8.24 ± 0.42 | 10.72 ± 0.05 | 0.29 | 11.45 ± 0.16 | 1.04 | 8.71 ± 0.53 |
| S189A | 9.01 ± 0.09 | -0.48 | 10.22 ± 0.16 | 0.73 | 7.93 ± 0.30 | 10.38 ± 0.07 | -0.03 | 11.21 ± 0.16 | 0.80 | 8.63 ± 0.44 |
| S202A | 5.92 ± 0.04 | -3.57 | 7.03 ± 0.10 | -2.46 | 8.45 ± 0.30 | 9.85 ± 0.06 | -0.56 | 10.61 ± 0.13 | 0.20 | 8.16 ± 0.25 |
| F288L | 8.88 ± 0.06 | -0.61 | 10.25 ± 0.10 | 0.76 | 7.90 ± 0.16 | 10.53 ± 0.06 | 0.12 | 11.27 ± 0.15 | 0.86 | 8.58 ± 0.42 |
| F313L | 8.11 ± 0.04 | -1.38 | 9.24 ± 0.08 | 0.25 | 8.14 ± 0.12 | 9.50 ± 0.06 | -0.91 | 10.29 ± 0.10 | -0.12 | 8.49 ± 0.27 |
| V317N | 7.40 ± 0.10 | -2.09 | 8.48 ± 0.22 | -1.01 | 8.11 ± 0.32 | 9.40 ± 0.13 | -1.01 | 10.06 ± 0.20 | -0.35 | 8.51 ± 0.48 |
| W321Y | 8.10 ± 0.09 | -1.39 | 9.51 ± 0.17 | 0.02 | 7.95 ± 0.21 | 8.76 ± 0.04 | -1.65 | 9.80 ± 0.12 | -0.61 | 8.38 ± 0.19 |

**Table S4.** Allosteric effect on WT and mutant DRD1 using G protein recruitment assay. MiniG_S_ recruitment using BRET assay and transfected HEKT cells in the presence of increasing concentrations of SKF81297 and LY3154207. Allosteric parameter (pK_B_) obtained by fitting dose response curves to “Allosteric EC50 shift” function of Graphpad Prism 8.4. Data are presented as mean values ± SEM with a minimum of two technical replicates and N = 3 biological replicates. **\***- denoted N = 2 biological replicates. Greek letter delta (Δ) for the difference (ΔpEC50) or affinity (ΔpK_B_) when compared with the wild-type receptor values. Data related to Fig. S9.

|  | SKF81297 |  | SKF81297/LY3154207 |  |  |  |
| --- | --- | --- | --- | --- | --- | --- |
|  | pEC<sub>50</sub> | ΔpEC<sub>50</sub> | pEC<sub>50</sub> | ΔpEC<sub>50</sub> | pK<sub>B</sub> | ΔpK<sub>B</sub> |
| WT | 8.79 ± 0.12 | 0.00 | 8.94 ± 0.06 | 0.00 | 8.04 ± 0.23 | 0.00 |
| W123A | 8.84 ± 0.10 | 0.05 | 8.95 ± 0.06 | 0.01 | 7.48 ± 0.10 | -0.56 |
| R130A | 8.67 ± 0.13 | -0.12 | 8.86 ± 0.05 | -0.08 | 6.08 ± 0.28 | -1.96 |
| A139L | 8.74 ± 0.13 | -0.05 | 8.75 ± 0.06 | -0.19 | 6.94 ± 0.27 | -1.10 |
| L143A | 8.67 ± 0.13 | -0.12 | 8.77 ± 0.06 | -0.17 | 7.90 ± 0.08 | -0.14 |
| W123A/A139L* | 8.64 ± 0.18 | -0.15 | 8.71 ± 0.07 | -0.23 | 5.68 ± 0.36 | -2.36 |

## METHODS

### Constructs design

The full-length gene coding sequence of human DRD1 was synthesized with no additional mutations or loop deletions added into the gene sequence. The N-terminal of DRD1 sequence was induced with FLAG tag followed by a fragment of β_2_AR N-terminal tail region (BN, hereafter) as fusion protein ^1^ while the C-terminal was attached with an 8×His tag to facilitate the protein expression and purification. The whole tag-attached DRD1 was then cloned into the standard pFastBac (Thermo Fisher) vector. The prolactin precursor sequence was inserted into the N terminus before the FLAG tag as signaling peptide to assist cell membrane localization and increase expression of DRD1. A miniG_αs_ format of G_αs_ with dominant negative mutations, G226A and A366S, was engineered (miniG_αs_DN_) according to the previous study ^2^. All the three G_s_ protein complex components, miniG_αs_DN_, rat G_β1_ and bovine G_γ2_, were constructed into pFastbac vector separately.

### Expression, assemble and purification of DRD1-G_s_ complexes

Before expression, the recombinant baculoviruses containing DRD1, miniG_αs_DN_, G_β1_ and G_γ2_ respectively were prepared using the Bac-to-Bac baculovirus expression system (Thermo Fisher). Cell cultures were grown to a density of 4×10^6^ cells/ mL in ESF 921 serum-free medium (Expression Systems). For the expression of the DRD1-G_s_ complexes, Sf9 cells were co-infected with the four types of baculoviruses prepared at 1:1:1:1 ratio. 48 hours after infection, the cultures were harvested by centrifugation at 1300×g (Thermo Fisher, H12000) for 20 min and frozen at −80°C for further usage.

For the purification of dopamine bound DRD1-miniG_s_ complexes, cell pellets from 1 L culture were thawed at room temperature and resuspended in buffer containing 20 mM HEPES pH 7.2, 75 mM NaCl, 5 mM CaCl_2_, 5 mM MgCl_2_, 10% Glycerol, 0.3 mM TECP, protease inhibitor cocktail (Bimake, 1 mL/ 100 mL suspension). The cell pellets were dounced to homogeneity and subsequently added with 1mM dopamine (MCE Inhibitors) to induce complexes formation on cell membrane. Half hour later, the cell suspension was further supplied with 10 μM LY3154207, treated with apyrase (25 mU mL^−1^, NEB) and incubated at room temperature for another one hour. Finally, the cell membrane in suspension was solubilized directly by addition of a detergent mixture containing 0.5% (w/v) dodecyl-β-D-maltoside (DDM, Anatrace), 0.1% (w/v) cholesteryl hemisuccinate TRIS salt (CHS, Anatrace), 0.025% (w/v) digitonin (Biosynth) and supplemented with 20 μM CID2886111 and 10 μg/mL Nb35. CID2886111 is another weaker PAM of DRD1 and bind to a divergent pocket in DRD1 compared to LY3154207. The membrane was solubilized for 3 hours at 4°C before separation by ultracentrifugation. The sample was centrifuged at 100,000 g (Ti45, Beckman) for 45 min. The isolated supernatant was then incubated for 2 hours at 4°C directly with FLAG resin (Smart-Lifesciences) pre-equilibrated with buffer containing 20 mM HEPES, pH 7.2, 100 mM NaCl. After batch binding, FLAG resin with immobilized protein complex was manually loaded onto a gravity flow column. Detergent was exchanged on FLAG resin by three washing steps in 20 mM HEPES, pH 7.2, 100 mM NaCl, 0.3 mM TCEP, 100 μM dopamine, 5 μM LY3154207, 10 μM CID2886111, supplied with different detergents: first 0.1% DDM, 0.02% CHS, 0.025% digitonin, then 0.02% DDM, 0.004% CHS, 0.05% digitonin, and finally 0.05% digitonin for 10 column volumes, each. The attached protein complex was then eluted in buffer containing 20 mM HEPES, pH 7.2, 100 mM NaCl, 0.3 mM TCEP, 100 μM dopamine, 5 μM LY3154207, 10 μM CID2886111, 0.05% digitonin, 200 μg/μL FLAG peptide.

For the purification of DRD1-miniG_s_ complex bound with both SKF81297 and PAM LY3154207, all the purification processes are the same as the above for dopamine bound complex with the following exceptions. The LY3154207 compound was added into the cell suspension at the concentration of 10μM after the first half-hour incubation with 10μM SKF81297 (Tocris) and together with the addition of apyrase (25 mU mL^−1^, NEB). No CID2886111 was supplied in the purification. In the later purification procedures, the buffer for washing and elution were all supplemented with 5 μM SKF81297 and 5 μM LY3154207.

Released protein was further concentrated to 0.5mL and then loaded onto a Superdex 200 10/300 GL increase column (GE Healthcare) pre-equilibrated with buffer containing 20 mM HEPES, pH 7.2, 100 mM NaCl, 0.05% digitonin, 0.1 mM TCEP, supplied with 100 μM dopamine, 5 μM LY3154207 and 10 μM CID2886111 or 5 μM SKF81297 and 5 μM LY3154207. The fractions of monomeric complex were pooled and concentrated for 20 folds (V/V), respectively. The final concentration of dopamine-LY3154207 bound D1R-G_s_ complex was about 10 mg mL^−1^ while concentration of SKF81297-LY3154207 bound D1R-G_s_ complex was 13 mg mL^−1^. Both concentrated protein samples were used for further electron microscopy experiments.

### Cryo-EM grid preparation and data collection

For the cryo-EM grid preparation, 3 μL of the purified DRD1-dopamine-LY3154207/ CID2886111-miniG_s_-Nb35 complex at the concentration about 10 mg mL^−1^, DRD1-SKF81297-LY3154207-miniG_s_-Nb35 complex at the concentration of 13 mg mL^−1^ were applied individually onto glow-discharged holey carbon grids (Quantifoil, Au200 R1.2/1.3) in a Vitrobot chamber (FEI Vitrobot Mark IV). The Vitrobot chamber was set to 100% humidity at 4 °C. Excess samples were blotted for 3 s and were vitrified by plunging into liquid ethane using a Vitrobot Mark IV (Thermo Fischer Scientific). Grids were stored in liquid nitrogen for screening and data collection usage.

For DRD1-dopamine-LY3154207/ CID2886111-miniG_s_-Nb35 complex, cryo-EM imaging was performed on a Titan Krios at 300kV in the Center of Cryo-Electron Microscopy, Zhejiang University (Hangzhou, China). Micrographs were recorded using a Gatan K2 Summit detector in counting mode with a pixel size of 1.014 Å using the SerialEM software ^3^. Movies were obtained at a dose rate of about 7.8 e/Å^2^/s with a defocus ranging from −0.5 to −3.0 μm. The exposure time was 8 s and 40 frames were recorded per micrograph. A total of 2002 movies were collected for the dopamine-LY3154207/CID2886111 bound DRD1-G_s_ complexes.

For the DRD1-SKF81297-LY3154207-miniG_s_-Nb35 complex, automatic data collection was performed on a FEI Titan Krios operated at 300kV in Cryo-Electron Microscopy Research Center, Shanghai Institute of Materia Medica, Chinese Academy of Sciences (Shanghai, China). The microscope was operated with a nominal magnification of 81,000× in counting mode, corresponding to a pixel size of 1.045 Å for the micrographs. A total of 3,607 movies for the dataset of DRD1-SKF81297-LY3154207-miniG_s_-Nb35 complex were collected by a Gatan K3 Summit direct electron detector with a Gatan energy filter (operated with a slit width of 20 eV) (GIF) using the SerialEM software ^3^. The images were recorded at a dose rate of about 26.7 e/Å2/s with a defocus ranging from −0.5 to −3.0 μm. The total exposure time was 3 s and intermediate frames were recorded in 0.083 s intervals, resulting in a total of 36 frames per micrograph.

### Image processing and map reconstruction

For the dopamine-bound DRD1-G_s_ complex, image stacks were aligned using MotionCor 2.1 ^4^. Contrast transfer function (CTF) parameters were estimated by Gctf v1.18 ^5^. The following data processing was performed using RELION-3.0-beta2 ^6^. Automated particle selection using Gaussian blob detection produced 1,432,690 particles. The particles were subjected to reference-free 2D classification to discard fuzzy particles, resulting in 679,229 particles for further processing. The map of GPBAR-G_s_ complex (EMD-30344) low-pass filtered to 60 Å was used as the reference map for 3D classification, generating one well-defined subset with 279,911 particles. Further 3D classifications focusing the alignment on the complex, produced three good subsets accounting for 212,256 particles, which were subsequently subjected to 3D refinement, CTF refinement and Bayesian polishing. The final refinement generated a map with an indicated global resolution of 3.2 Å at a Fourier shell correlation of 0.143.

For DRD1-SKF81297-LY3154207-miniG_s_-Nb35 complex, movie stacks were subjected to beam-induced motion correction using MotionCor2.1 ^4^. Contrast transfer function parameters for each micrograph were determined by Ctffind4 ^7^. The micrographs with the measured resolution worse than 4.0 Å and micrographs imaged within carbon area were discarded, generating 2,842 micrographs for further processing. Particle selection, 2D and 3D classifications were performed on a binned dataset with a pixel size of 2.09 Å using RELION-3.0-beta2 ^6^. About 2000 particles were manually selected and subjected to 2D classification. Representative averages were picked as template for auto-picking. The auto-picking process produced 2,193,368 particles, which were subjected to reference-free 2D classifications to exclude fuzzy particles. The 3D density map of DRD1-SKF83959-G_s_ ^2^ low-pass filtered to 40 Å was served as initial reference map for seven rounds 3D classifications, resulting in a single well-defined subset with 249,418 particles. Subsequent 3D refinement, CTF refinement, Bayesian polishing and DeepEnhancer ^8^ processing generated a map with an indicated global resolution of 3.0 Å at a Fourier shell correlation of 0.143.

### Structure model building and refinement

The structure of DRD1-SK81297-G_s_ (PDB: 7JV5) was used as initial model for model rebuilding and refinement against the electron microscopy maps of DRD1-G_s_ complexes. The PDB models of LY3154207, and dopamine were generated with ChemSketch (https://www.acdlabs.com/resources/freeware/chemsketch/index.php). The initial models were docked into the electron microscopy density maps using Chimera ^9^ followed by iterative manual adjustment and rebuilding in COOT ^10^. Real space refinement and reciprocal space refinement were performed using Phenix programs ^11^. The model statistics were validated using MolProbity ^12^. Structure figures were prepared in Chimera and PyMOL (https://pymol.org/2/). The final refinement statistics are provided in Tables S1.

### G_s_ Bioluminescence Resonance Energy Transfer (BRET) Recruitment Assays

To measure G protein recruitment (BRET), HEK293T cells (ATCC CRL-11268) maintained in DMEM containing 10% (v/v) dialyzed FBS, 1 IU ml^−1^ Penicillin G, and 100 μg mL^−1^ Streptomycin were passed to 10 cm dishes and co-transfected using TransIT (Mirus Bio) in an approximate 1:2.5 ratio with DRD1 containing C-terminal *Renilla* luciferase (*R*Luc) and Venus-tagged N-terminal MiniG protein ^13^. After at least 24 hours, transfected cells were plated in poly-lysine coated 96-well white clear bottom cell culture plates in plating media (DMEM containing 1% (v/v) dialyzed FBS, 1 IU mL^−1^ Penicillin G, and 100 μg ml^−1^ Streptomycin) at a density of 40,000 cells in 200 μL per well and incubated overnight.

The following day, media was aspirated and cells were washed once with 60 μL of drug buffer (1X HBSS, 20 mM HEPES, pH 7.4). Then 60 μL of drug buffer was added per well and drug stimulation was performed with the addition of 15 μL of 6× drug dilution of LY3154207 in drug dilution buffer (1× HBSS, 20 mM HEPES, 0.3% (w/v) BSA, 0.03% (w/v) ascorbic acid, pH 7.4) per well and incubated at RT. After 60 minutes of incubation, 10 μL of the *R*Luc substrate, coelenterazine h (Promega) at 5 μM final concentration was added per well. After an additional 5 minutes, 15 μL of 6× drug dilution of SKF81297 in drug dilution buffer was added to the each well, plates were read for both luminescence at 485 nm and fluorescent eYFP emission at 530 nm for 1 second per well using a Mithras LB940 (Berthold Technologies). Plates were read for multiple time points up to 30 minutes. The BRET ratio of eYFP/*R*Luc was calculated per well and the net BRET ratio was calculated by subtracting the eYFP/*R*Luc ratio per well from the eYFP/*R*Luc ratio in wells without Venus-tagged N-terminal MiniG protein present. The net BRET ratio was normalized to the no drug addition and plotted as a function of drug concentration using Graphpad Prism 8 (Graphpad Software Inc., San Diego, CA).

### DRD1 G_s_-mediated G_s_-cAMP Accumulation Assay

DRD1 G_s_-mediated G_s_-cAMP accumulation assays with HEK293T (ATCC CRL-11268) were performed using cells transiently expressing human DRD1 and the cAMP biosensor GloSensor-22F (Promega). Cells were seeded (20,000 cells/35 μL/well) into white 384 clear-bottom, tissue culture plates in DMEM containing 1% (v/v) dialyzed fetal bovine serum (FBS). Next day, 3× drug dilutions were diluted in HBSS, 20 mM N-(2-hydroxyethyl) piperazine-N’-ethanesulfonic acid (HEPES), 0.3% (w/v) bovine serum albumin (BSA), 0.03% (w/v) ascorbic acid, pH 7.4. Media was decanted from 384 well plates and 20 μL of drug buffer (HBSS, 20 mM HEPES, pH 7.4) containing GloSensor reagent was added per well and allowed to equilibrate for at least 15 min at room temperature. Cells were then treated with 10 μL per well of 3× drug using a FLIPR (Molecular Devices). After 15 min, G_s_-cAMP accumulation was read on a TriLux Microbeta (PerkinElmer) plate counter. Data were normalized to maximum G_s_-cAMP accumulation by dopamine (100%). Data were analyzed using the sigmoidal log(agonist) vs. dose response function built into GraphPad Prism 8.4.

### Figure preparation

The density maps were prepared in UCSF Chimera (https://www.cgl.ucsf.edu/chimera/) and and UCSF ChimeraX (https://www.cgl.ucsf.edu/chimerax/). Structural comparisons and alignments figures were prepared with PyMOL (https://pymol.org/2/).

